# A deep learning approach to identify new gene targets of a novel therapeutic for human splicing disorders

**DOI:** 10.1101/2020.02.03.932103

**Authors:** Dadi Gao, Elisabetta Morini, Monica Salani, Aram J. Krauson, Ashok Ragavendran, Serkan Erdin, Emily M. Logan, Anil Chekuri, Wencheng Li, Amal Dakka, Nikolai Naryshkin, Chris Trotta, Kerstin A. Effenberger, Matt Woll, Vijayalakshmi Gabbeta, Gary Karp, Yong Yu, Graham Johnson, William D. Paquette, Michael E. Talkowski, Susan A. Slaugenhaupt

**Affiliations:** Center for Genomic Medicine, Massachusetts General Hospital Research Institute, Boston, MA; Department of Neurology, Massachusetts General Hospital Research Institute and Harvard Medical School, Boston, MA; Program in Medical and Population Genetics and Stanley Center for Psychiatric Research, Broad Institute of Harvard and MIT, Cambridge, MA; PTC Therapeutics, Inc., South Plainfield, NJ; NuPharmAdvise LLC, Sanbornton, NH; Albany Molecular Research Inc., Albany, NY

## Abstract

Pre-mRNA splicing is a key control point in human gene expression. Disturbances in splicing due to mutation or aberrant splicing regulatory networks lead to dysregulated protein expression and contribute to a substantial fraction of human disease. Several classes of active and selective splicing modulator compounds have been recently identified, thus proving that pre-mRNA splicing is a viable target for therapy. We describe herein the identification of BPN-15477, a novel splicing modulator compound, that restores correct splicing of exon 20 in the Elongator complex protein 1 *(ELP1)* gene carrying the major IVS20+6T>C mutation responsible for familial dysautonomia. We then developed a machine learning approach to evaluate the therapeutic potential of BPN-15477 to correct splicing in other human genetic diseases. Using transcriptome sequencing from compound-treated fibroblast cells, we identified treatment responsive sequence signatures, the majority of which center at the 5’ splice site of exons whose inclusion or exclusion is modulated by SMC treatment. We then leveraged this model to identify 155 human disease genes that harbor ClinVar mutations predicted to alter pre-mRNA splicing as potential targets for BPN-15477 treatment. Using *in vitro* splicing assays, we validated representative predictions by demonstrating successful correction of splicing defects caused by mutations in genes responsible for cystic fibrosis (*CFTR*), cholesterol ester storage disease (*LIPA*), Lynch syndrome (*MLH1*) and familial frontotemporal dementia (*MAPT*). Our study shows that deep learning techniques can identify a complex set of sequence signatures and predict response to pharmacological modulation, strongly supporting the use of *in silico* approaches to expand the therapeutic potential of drugs that modulate splicing.

---

RNA splicing is a complex and tightly regulated process that removes introns from pre-mRNA transcripts to generate mature mRNA. Differential processing of pre-mRNA is one of the principal mechanisms generating diversity in different cell and tissue types. This process can give rise to functionally different proteins and can also generate mRNAs with different localization, stability and efficiency of translation through alternative splicing of UTRs. RNA splicing requires the widely conserved spliceosome machinery along with multiple splicing factors^1^. The splicing reaction is directed by specific sequences, including the 5’ and 3’ splice sites, the intron branch point, and splice site enhancers and silencers found in both exons and introns^2^. Changes in the sequence of these elements, through inherited or sporadic mutations, can result in deficient or aberrant splice site recognition by the spliceosome. Disruption of splicing regulatory elements can generate aberrant transcripts through complete or partial exon skipping, intron inclusion or mis-regulation of alternative splicing, while mutations in the UTRs may affect transcript localization, stability or efficiency of translation. Mutations that alter mRNA splicing are known to lead to many human monogenic diseases including spinal muscular atrophy (SMA), neurofibromatosis type 1 (NF1), cystic fibrosis (CF), familial dysautonomia (FD), Duchenne muscular dystrophy (DMD) and myotonic dystrophy (DM), as well as contribute to complex diseases such as cancer and diabetes^3-18^. The emergence of high throughput sequencing of large disease cohorts^19-21^, and the remarkable efforts to aggregate and annotate these mutations in an accessible infrastructure such as ClinVar^22^, now provides an unprecedented opportunity to apply novel deep learning approaches to predict mutations that affect pre-mRNA splicing^23^. The potential of developing such models will continue to increase as next generation transcriptome sequencing (RNASeq) data are amassed and curation of the associated mutational processes matures ^23-26^.

New therapeutic approaches aimed at correction of pre-mRNA splicing defects, including antisense oligonucleotides, splicing modulator compounds (SMCs), and modified exon-specific U1 small nuclear RNA, have shown significant promise in many diseases^27-35^. SMCs are attractive because they can be optimized for broad tissue distribution and are orally administered^27,36,37^. With advances in precision medicine and the capability to discover patient-specific mutations, there is strong impetus to develop new methods to predict if a drug might be beneficial in a specific patient. Deep learning techniques offer the potential to accomplish this at scale by integrating genomic data with annotation databases and relevant information about mutational mechanisms^38^. Deep learning models have been successfully applied to a spectrum of biological topics, including genotype-phenotype correlation studies^39^, identification of disease biomarkers^40^, and identification of protein binding motifs^41^. Here, we applied and optimized a specific deep convolutional neural network (CNN) to discover motifs that that are likely to be responsive to BPN-15477, a potent SMC of the *ELP1* pre-mRNA carrying the major FD splice mutation IVS20+6T>C. We identified 155 genes harboring pathogenic ClinVar mutations, each predicted to disrupt pre-mRNA splicing, that could be corrected by BPN-15477 treatment, and validated several using minigenes or patient cells. These studies suggest that the integration of genomic information, clinical annotation of disease associated variants, and deep learning techniques have significant potential to predict therapeutic targeting for precision medicine.

## Results

### Discovery of the splicing modulator BPN-15477

Several small-molecules have been developed to selectively modulate the splicing of specific pre-mRNAs, offering potential new treatments for SMA and FD^27,36,37,42-45^. One such compound, kinetin (6-furfurylaminopurine), was previously shown to promote exon 20 inclusion in the Elongator complex protein 1 gene (*ELP1*, MIM: 603722) in FD^44,46^. Although kinetin is a naturally occurring compound with a safe absorption, distribution, metabolism, and excretion (ADME) profile, very high doses are necessary to achieve modest *ELP1* splicing correction *in vivo* ^47,48^. As part of the NIH Blueprint Neurotherapeutics Network we identified a new class of highly potent SMCs that selectively modulate ELP1 pre-mRNA splicing and increase the inclusion of exon 20^49^. Through further optimization we identified BPN-15477 (Fig.1a), a compound that increases full-length *ELP1* mRNA by increasing exon 20 inclusion. BPN-15477 is significantly more potent and efficacious than kinetin in our luciferase splicing assay (Fig.1b) and it rescues ELP1 splicing in FD patient cell lines (Fig.1c).

**Fig. 1.**
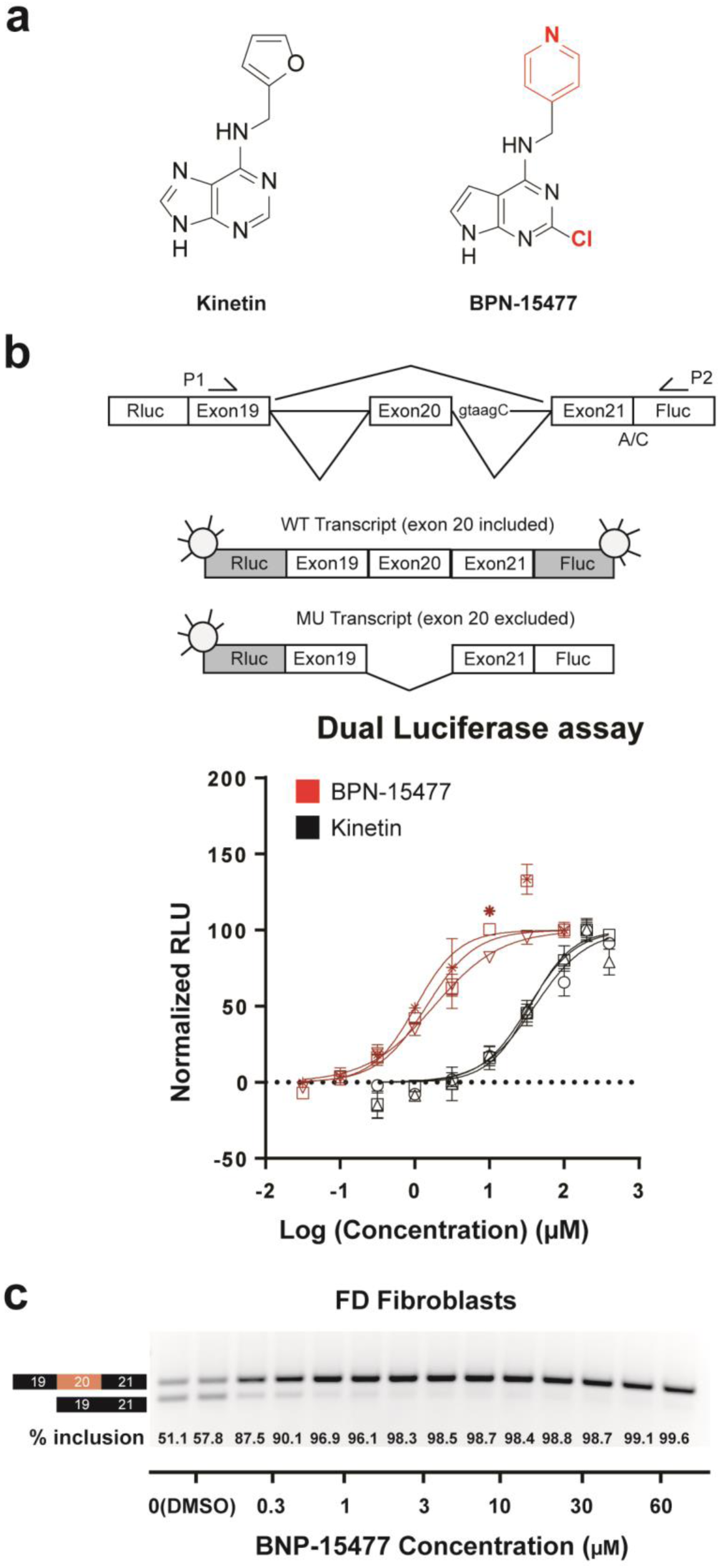
Identification of the small molecule splicing modulator BPN-15477. (**a**) Molecular structure of BPN-15477 compared to kinetin, the northern heterocycle and C-2 substitution are indicated in red. (**b**) Schematic representation of the dual-reporter minigene used to measure splicing. Rluc and Fluc indicate Renilla and Firefly luciferase, respectively. A/C indicates the start codon mutation in Fluc and taagC indicates the location of the FD mutation. Dose response curves for kinetin and BPN-15477. Rluc-FD-Fluc transfected HEK293T cells treated for 24 hours. Normalized relative luciferase units (RLU), which measure exon 20 inclusion, are plotted as a function of compound concentration. Assays were run in triplicate and curves were created by nonlinear regression using Prism4 (GraphPad Software Inc.). **(c)** Validation of BPN-15477 splicing correction in FD fibroblasts. Cells were treated in duplicate for 24 hours at the concentrations indicated.

### Evaluation of BPN-15477 on transcriptome splicing

To estimate the influence of BPN-15477 on transcriptome-wide splicing, we treated six wildtype (WT) human fibroblast cell lines with 30 μM BPN-15477 or vehicle (DMSO) for seven days and performed RNA-seq to evaluate changes in exon inclusion (Supplementary Table 1). Splicing differences were determined by counting the RNASeq reads covering two junctions of three consecutive exons (exon triplets) and by comparing the change in percent spliced-in (ΔPSI or Δψ) of the middle exon after treatment (Fig. 2a, see Methods)^50,51^. We identified 934 exon triplets that showed differential middle exon inclusion or exclusion in response to BPN-15477 treatment; 254 with increased exon inclusion (Δψ ≥ 0.1 and FDR < 0.1), and 680 with increased exon exclusion (Δψ ≤ −0.1 and FDR < 0.1, Fig 2b and Supplementary Table 2). BPN-15477 modulates splicing selectively as we observed splicing changes in only 0.58% of all expressed triplets (934 out of 161,097 expressed triplets). To experimentally confirm the accuracy of the PSI changes measured by RNASeq, we performed independent treatment experiments and evaluated exon inclusion using RT-PCR (Fig. 2d-g) and found the estimated percent exon inclusion to be remarkably consistent with the calculated Δψ values (Fig. 2c).

**Fig. 2.**
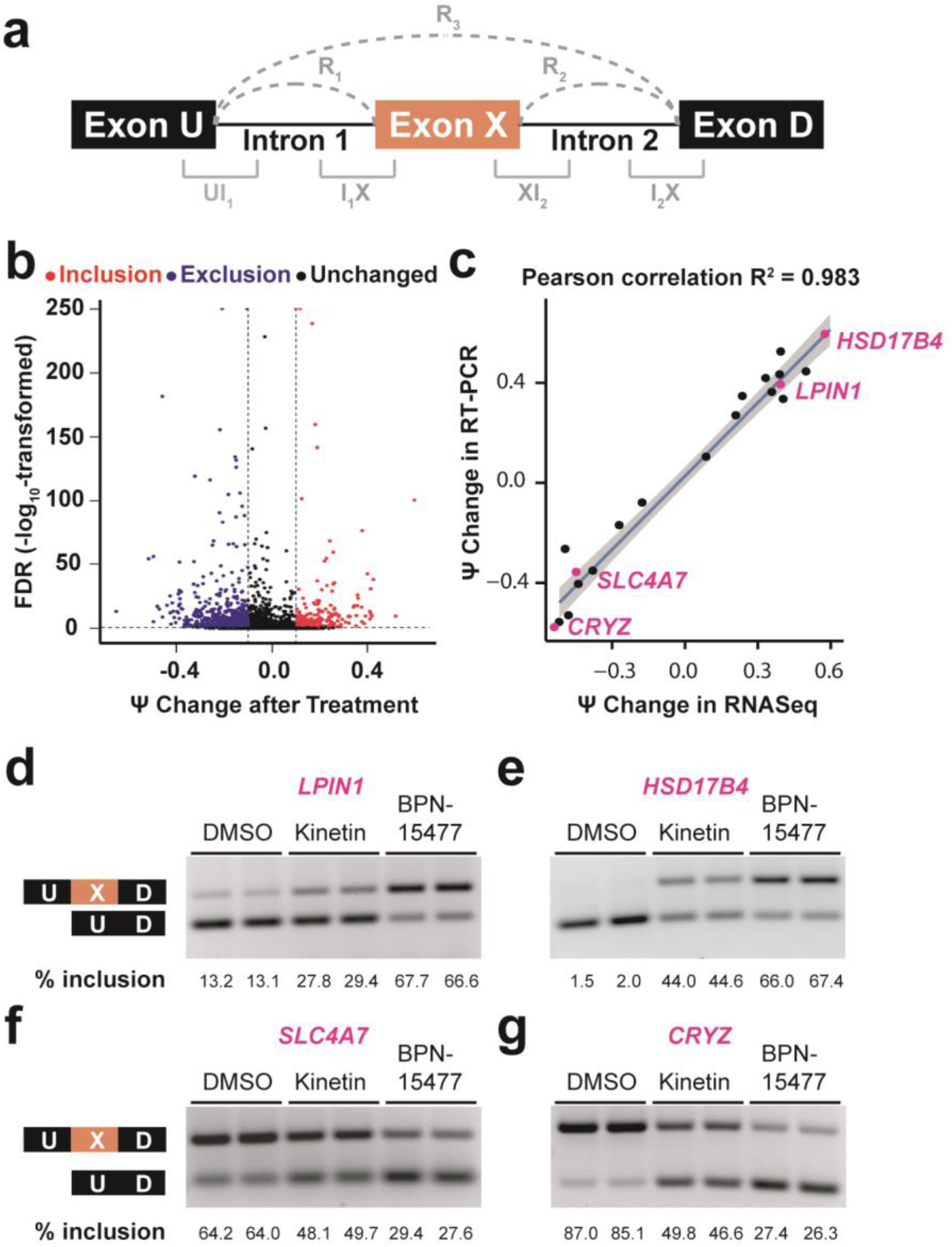
Transcriptome-wide changes in response to BPN-15477. **(a)** Schematic representation of an exon triplet. R_1_, R_2_ and R_3_ represent RNASeq reads spanning two adjacent exons. The region marked UI_1_, I_1_X, XI_2_ and I_2_D represent 25 exonic base pairs and 75 intronic base pairs. **(b)** Volcano plot showing the ψ changes after treatment with BPN-15477. Each dot represents one of the 161,097 expressed exon triplets in human fibroblasts. The x axis represents the ψ change after the treatment and the y axis represents the false discovery rate (FDR) (log10 transformed). Red dots represent the exon-triplets with an increase of middle exon inclusion (Δψ ≥ 0.1 and FDR < 0.1) while blue dots represent an increase of middle exon exclusion (Δψ ≤ −0.1 and FDR < 0.1). Black dots are exon-triplets not responsive to the treatment. The two vertical dashed lines indicate a ψ change of 0.1 and −0.1 as thresholds for exclusion and inclusion, respectively. The horizontal dashed line indicates an FDR of 0.1. **(c)** Independent RT-PCR validation of splicing changes of twenty randomly selected candidates. For each validated exon-triplet, ψ change measured by RNASeq (x axis) is plotted against the splicing changes measured by RT-PCR (y axis). The R^2^ value indicates the coefficient of Pearson correlation. The solid line shows the estimated linear regression. The grey zone indicates the 95% confidence interval for predictions from the estimated linear regression. **(d-g)**, Representative RT-PCR validations of splicing response of *LPIN1 (d), HSD17B4 (e), SLC4A7(f)* and *CRYZ (g)* in treated human fibroblasts. The upper bands indicate the isoform in which the middle exon is included while the lower bands indicate the isoform in which the middle exon is skipped. As expected, BPN-15477 is more potent than kinetin.

### Convolutional neural network identifies sequence signatures responsible for BPN-15477 response

Pre-mRNA splicing is regulated by both exonic and intronic sequence elements. These sequences govern interaction with the spliceosome and splicing factors and regulate the fate of exon recognition and inclusion^2,52^. We hypothesized that sequence signatures within the exon triplets are a key determinant of drug responsiveness. To identify the sequence motifs, we trained a CNN model using the inclusion-response set (254 exon triplets), exclusion-response set (680 exon triplets) and the unchanged-response set (382 exon triplets with two expressed isoforms, Δψ < 0.01 and FDR ≥ 0.1, Supplementary Table 2). The network consisted of two layers of convolutions with a total of 2.5 million trainable parameters (Supplementary Fig. 1a) and was optimized for predicting splicing changes in every given exon triplet after BPN-15477 treatment.

Our model achieved an average area under the curve (AUC) of 0.85 and identified thirty-nine 5-mer motifs that best explain drug responsiveness (Supplementary Figs. 1b-c and 2). Twelve of these motifs explained 94.87% of the AUC, each of which altered more than 0.1 of AUC for at least one class of prediction (Fig. 3a). Importantly, analysis of these motifs revealed that proximity to, and in many cases as part of, the 5’ splice site of the middle exon had the largest influence in modulating treatment response (Fig. 3b, Supplementary Fig. 3a). To assess the robustness of the motifs identified by our CNN model, we compared the similarities of the twelve CNN-identified motifs with the most enriched five-nucleotide combinations (5-mer) at 5’ splice sites of the middle exons in our training set. The 5-mer sequence enrichment analysis for nucleotides at positions −3 to +7 of 5’ splice sites agreed with motifs identified by the CNN model (Supplementary Fig. 3b). Examination of our predicted motifs, combined with the 5-mer and positional analysis, shows that the 5’ splice sites of responsive exons are non-canonical. Therefore, we evaluated the strength of the four triplet splice sites using MaxEntScan^53^ and found that the 5’ splice site of the middle exon is significantly weaker when exon inclusion is enhanced by treatment (Fig. 3c). These results are consistent with previous findings that support the role of this class of SMCs in promoting the recruitment of U1 snRNP to non-canonical 5’ splice sites^45,46^.

**Fig. 3.**
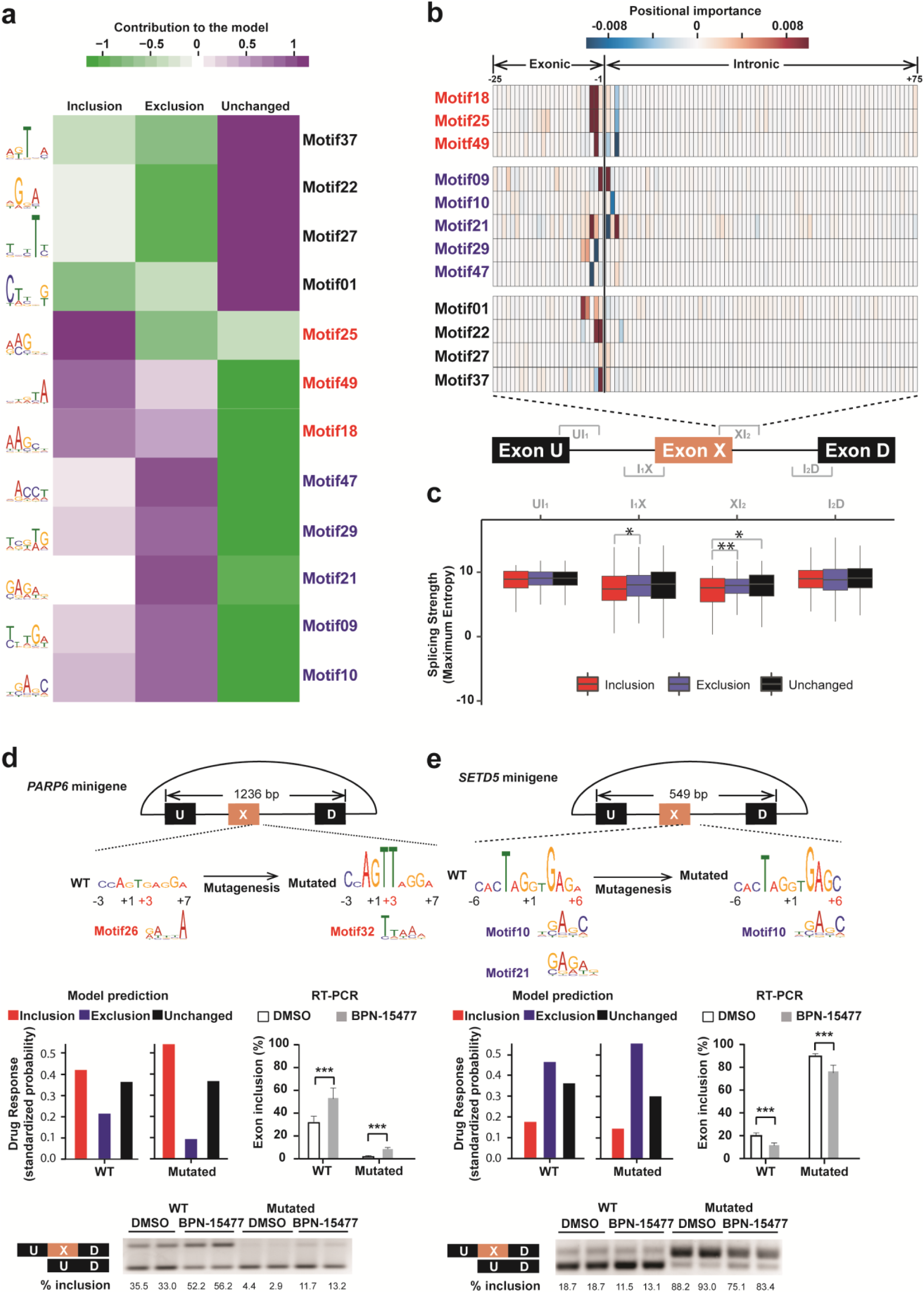
Convolutional neural network achieves high performance in predicting drug response and reveals sequence motifs that contribute to drug sensitivity. **(a)** Heatmap of top 12 motifs predicted to contribute to inclusion (red), exclusion (blue) or unchanged (black) response. The color bar indicates the directional contribution of each motif. The purple domain indicates positive contribution while the green domain indicates negative contribution. The LOGO plot of each motif is shown on the left side of the heatmap. **(b)** Heatmap of motif importance at each position within XI_2_ region at the 5’ splice site of the middle exon. The thick vertical line shows the exon-intron boundary. The bar indicates the positional importance. **(c)** Box plot showing the strength of each splice junction along the exon triplets for inclusion (red), exclusion (blue) and unchanged (black) group defined by the RNASeq. The middle lines inside boxes indicate the medians. The lower and upper hinges correspond to the first and third quartiles. Each box extends to 1.5 times inter-quartile range (IQR) from upper and lower hinges, respectively. Outliers are not shown. Only comparisons with significant difference are marked by stars (two-tailed, unpaired Welch’s *t* test with Bonferroni correction). **(d-e)**, *Upper row*: Schematic representations of *PARP6* and *SETD5* triplet minigene. The length of the exon triplets cloned into the minigenes are shown. The sequences adjacent to the 5’ splice site of the middle exon are shown in LOGO plots. The height of each nucleotide was estimated using *in silico* saturated mutagenesis (See Methods). The red coordinate numbers indicate the positions of mutations relative to the 5’ splice sites. Their closely matched CNN motifs are indicated beneath. *Middle row*: Splicing changes of the middle exons in both wildtype and mutated exon triplets, predicted by the CNN model (*left*) and measured by RT-PCR of the minigene (*right*). Each RT-PCR was done in duplicate and independently repeated three times for each minigene to make the bar plots (two-tailed, unpaired Student’s *t* test). *Bottom row:* Example of splicing changes induced by the treatment using a minigene splicing assay. The percentage of middle exon inclusion is indicated beneath each lane. The statistical significance is determined via two-tailed, unpaired Student’s test. In the figure, \**p* < 0.05; \*\**p* < 0.01; \*\*\**p*<0.001.

We next sought to determine if our CNN model could predict treatment response in mutated exon triplets. We generated minigenes for treatment responsive alternatively spliced triplets in *PARP6, SETD5* and *CPSF7*. These genes were chosen because their genomic triplet length enabled evaluation in an appropriate splicing vector. We mutated nucleotides in silico between positions +2 to +6 in the 5’ splice site of each triplet and used our CNN model to predict treatment response. Mutations predicted to be responsive were introduced into the minigenes and splicing evaluated by RT-PCR. In all three minigene splicing assays, the RT-PCR data confirmed our CNN model predictions for mutant triplets (Fig. 3d-e, Supplementary Fig. 3c). Therefore, our CNN model will allow us to evaluate the potential therapeutic value of BPN-15477 on human splicing mutations.

### Identification of potential therapeutic targets of BPN-15477

To evaluate the predictive power of our CNN model to determine which human disease-causing mutations might respond to treatment with BPN-15477, we first identified the pathogenic mutations that alter splicing in ClinVar. We considered all 89,642 annotated pathogenic or likely-pathogenic mutations (CV-pMUTs) and predicted their influence on splicing using SpliceAI^23^. We found that ∼20% of all CV-pMUTs are predicted to alter splicing within 5 kb of the mutation, and that ∼80% of these disrupt Ensembl-annotated splice sites (GRCh37 version 75). As expected, the CV-pMUTs disrupting annotated splice sites were substantially closer to the splicing junction than the overall CV-pMUTs (Fig. 4b); ∼98% of them were within 75bp. This observation confirms that our 100 bp training regions UI_1_, I_1_X, XI_2_ and I_2_D captured the majority of the potentially targetable pathogenic splicing alterations. We next used our CNN model to predict which genes harboring CV-pMUTs might respond to BPN-15477 treatment. The 14,272 CV-pMUTs that disrupt annotated splice sites are mapped to 11,616 exon triplets, and our CNN model predicted that 271 of these triplets should be responsive to BPN-15477 treatment (Supplementary Table 3). Therefore, our analysis identified 155 genes that harbor 214 annotated disease-causing mutations that could be targets for splicing correction using BPN-15477. The responsive genes containing the top 20 most frequent mutations and their associated human diseases are shown in Table 1 (gnomAD v2.1.1). Examination of Table 1 demonstrates the remarkable therapeutic potential of this class of splicing modulators.

**Table 1.** ClinVar pathogenic mutations predicted to be rescued by BPN-15477 treatment and selected based on top populational allele frequencies in gnomAD (v2.1.1)

| Gene | Mutation | Frequency | Molecular consequence (ClinVar) | Splicing (SpliceAI) | Predicted drug response | Disease |
| --- | --- | --- | --- | --- | --- | --- |
| *IL36RN* | c.115+6T>C | 0.00095504 | Intronic | Loss | Inclusion | Pustular psoriasis |
| *CA5A* | c.555G>A | 0.00016644 | Synonymous | Loss | Inclusion | Carbonic anhydrase VA deficiency |
| *ORC6* | c.449+5G>A | 0.00016296 | Intronic | Loss | Inclusion | Meier-Gorlin syndrome 3 |
| *DNAH9* | c.1970+4A>G | 0.00014584 | Intronic | Loss | Inclusion | Ciliary dyskinesia |
| *HBB* | c.92G>A | 1.03E-04 | Missense | Loss | Inclusion | Beta thalassemia |
| *LIPA* | c.894G>A | 0.00010262 | Synonymous | Loss | Inclusion | Lysosomal acid lipase deficiency |
| *PIGN* | c.963G>A | 8.03E-05 | Synonymous | Loss | Inclusion | Multiple congenital anomalies-hypotonia-seizures syndrome 1 |
| *NSD1* | c.6152-5T>G | 5.96E-05 | Intronic | Loss | Inclusion | Beckwith-Wiedemann syndrome |
| *DGUOK* | c.591G>A | 4.95E-05 | Intronic / Synonymous | Loss | Inclusion | Mitochondrial DNA-depletion syndrome 3 |
| *SRD5A2* | c.547G>A | 4.88E-05 | Missense | Loss | Inclusion | 3-Oxo-5 alpha-steroid delta 4-dehydrogenase deficiency |
| *GLA* | c.639+919G>A | 4.53E-05 | Intronic | Gain | Exclusion | Fabry disease |
| *ATM* | c.2250G>A | 4.39E-05 | Synonymous | Loss | Inclusion | Ataxia-telangiectasia syndrome |
| *PARN* | c.659+4_659+7 delAGTA | 4.38E-05 | Intronic | Loss | Inclusion | Dyskeratosis congenita |
| *POLG* | c.3104+3A>T | 3.98E-05 | Intronic | Loss | Inclusion | Progressive sclerosing poliodystrophy |
| *GYP A* | c.232G>A | 3.60E-05 | Missense | Loss | Inclusion | Blood group erik |
| *GPX4* | c.476+5G>A | 3.59E-05 | Intronic | Loss | Inclusion | Spondylometaphyseal dysplasia |
| *BRCA1* | c.4675G>A | 3.19E-05 | Missense | Loss | Inclusion | Breast-ovarian cancer, familial 1 |
| *SPTA1* | c.6600+5G>T | 3.19E-05 | Intronic | Loss | Inclusion | Congenital hemolytic anemia |
| *TPO* | c.1768G>A | 3.18E-05 | Missense / Intronic | Loss | Inclusion | Deficiency of iodide peroxidase |
| *ACADSB* | c.922G>A | 2.95E-05 | Missense | Loss | Inclusion | Deficiency of 2-methylbutyryl-CoA dehydrogenase |

**Fig. 4.**
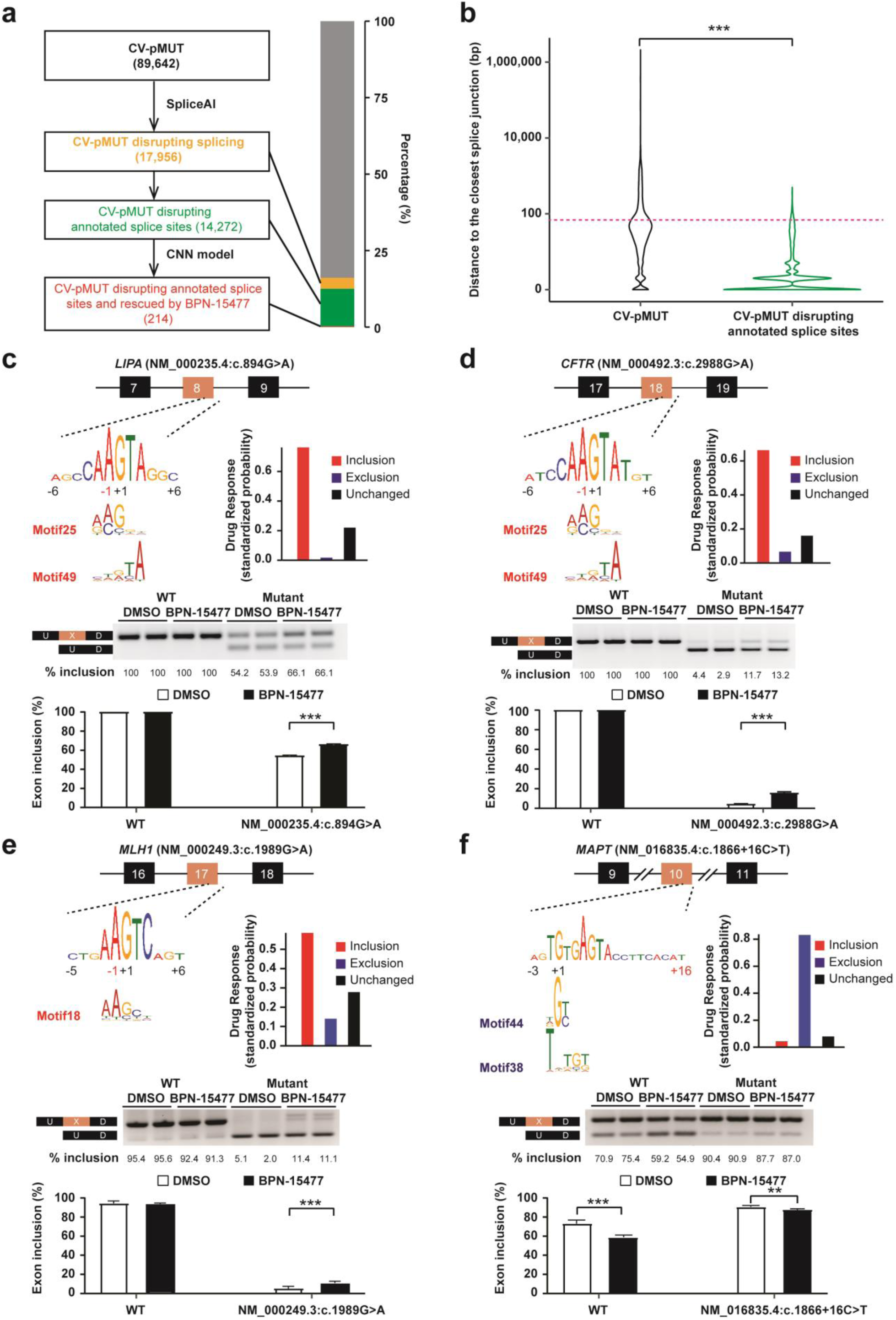
Identification of new therapeutic targets for BPN-15477. **(a)** Workflow to identify all potential therapeutic targets for BPN-15477. SpliceAI was applied on all ClinVar pathogenic mutations (CV-pMUTs) and the CNN model was used to determine whether counts per million (CPMs) disrupting annotated splice sites would be rescued by BPN-15477 treatment (*left*). The bar plot shows the percentage of each. **(b)** Violin plot of the distance of all CV-pMUTs or the CV-pMUTs disrupting annotated splicing to the closest splice junction. Y axis is a log_10_-transformed scale. Each violin shape shows the distribution of distance. The pink horizontal dashed line indicates 75 bp from the closest splice junction. The significance of difference is determined via Kolmogorov-Smirnov (K-S) test. **(c-f)**, *Upper row*: The sequences at the 5’ splice site of *LIPA* Exon 8 in patient cells (c) and minigene constructs for *CFTR, MLH1* and *MAPT* (d-f) are shown. The sequences around the 5’ splice sites of the middle exon are shown in LOGO plots, with closely matched CNN motifs indicated below. The red coordinate numbers indicate the position of mutations introduced mutations. The bar plots demonstrate the CNN model prediction of treatment response for the mutated sequences. *Middle row*: RT-PCR validation of treatment responses in cell lines carrying the specific splice mutation or the mutated minigene. Each set of gels is one of the triplicates used to generate the bar plots beneath. *Bottom row*: The bar plots demonstrate the splicing change promoted by BPN-15477 treatment. The statistical significance is determined via two-tailed, unpaired Student’s test. In the figure, \**p* < 0.05; \*\**p* < 0.01; \*\*\**p*<0.001.

### Experimental validation of new therapeutic targets for BPN-15477

To evaluate whether BPN-15477 ameliorates aberrant splicing events, we searched for available human cell lines carrying the splicing mutations predicted to respond to our treatment (Supplementary Table 3). From the Coriell cell repository we were able to obtain a cell line with the c.894G>A mutation in the *LIPA* gene (Table 1)^54^. Mutations in *LIPA* cause both the severe infantile-onset Wolman disease and the milder late-onset cholesterol ester storage disease (CESD)^55-57^ (MIM# 278000)^58^. The c.894G>A mutation leads to skipping of exon 8 and is responsible for the milder CESD. This is the most common *LIPA* gene mutation and it is found in about half of individuals with LAL deficiency^59^. To validate the treatment effect on the splicing of exon 8, patient cells were treated with 60 µM of BPN-15477 for 24 hours. As predicted, the treatment promoted the inclusion of exon 8, with mutated cells showing a 10% increase in normal transcript levels, which would lead to increased functional lipase A protein in patients (Fig. 4c 3). To overcome the limitations in available patient cell lines harboring specific splicing mutations, we prioritized our experimental validations based on the availability of minigene constructs. We obtained HEK293 cells stably expressing a minigene containing the full length *CFTR* coding sequence carrying a c.2988G>A mutation and flanking introns (Supplementary table 3)^60^. This mutation is associated with abnormal CFTR function and causes a mild form of CF (MIM# 219700)^61^. Treatment with BPN-15477 increased exon 18 inclusion by 10%, confirming our CNN model prediction (Fig. 4d). This is exciting since a small increase in functional CFTR protein is associated with a significant improvement in patient phenotype. We also generated a *MLH1* minigene spanning exons 16 to 18 harboring the c.1989 G>A mutation (Supplementary table 3), which leads to hereditary nonpolyposis colorectal cancer (HNPCC) or Lynch syndrome (MIM# 120435) and to skipping of exon 17 ^14,62^. As predicted, the treatment significantly increased exon 17 inclusion in cell lines expressing the mutated minigene (Fig 4e, Supplementary Table 3).

Finally, to validate the utility of our treatment to promote exon skipping as a therapeutic modality, we searched for drug responsive disease-causing mutations that lead to abnormal middle exon inclusion. The MAPT gene is associated with familial frontotemporal dementia and parkinsonism linked to chromosome 17 (FTDP-17, MIM# 600274)^63-66^. In healthy human brains, the alternative splicing of *MAPT* exon 10 is strictly regulated to maintain equal amounts of the 3R (exon 10 skipped) and 4R (exon 10 retained) tau isoforms. Disruption of this balance by increased 4R tau increases the risk of developing FTD. The c.1866 +3 G>A mutation, commonly referred to IVS10 +3, in the *MAPT* gene increases the inclusion of exon 10 and is associated with FTDP-17. Our CNN model predicted that the mutated sequence would be responsive to BPN-15477 and the treatment would promote exon 10 exclusion, thereby potentially restoring 3R/4R balance (Supplementary Table 3). We generated HEK293 cells stably expressing a *MAPT* minigene encompassing exons 9 to 11 and partial introns 9 and 10 with both the wild type sequence and the c.1866 +3 G>A mutation. As predicted, treatment with BPN-15477 led to a significant reduction of exon 10 inclusion in the wild type MAPT transcript. Unfortunately, introduction of the c.1866 +3 G>A mutation in the minigene context completely disrupted exon 10 skipping in both the DMSO and BPN-15477 treated minigenes (Supplementary Fig. 4). However, it is known that in the human brain the c.1866 +3G>A mutation “leaks” a detectable amount of 3R tau isoform^67^, suggesting that treatment with a splicing modulator *in vivo* might still increase the level of 3R tau.

Given the importance of identifying therapeutics that modulate the 3R/4R tau ratio, combined with our demonstration that BPN-15477 can promote skipping of MAPT exon 10, we next evaluated the most common MAPT mutation, a C to T substitution in intron 10, c.1866+16C>T, commonly called IVS10 +16. Although this mutation increases inclusion of exon 10^63^, it did not appear in our list of responsive mutations because SpliceAI did not correctly predict its effect on splicing. We applied our CNN model to the MAPT c.1866+16C>T exon triplet, which predicted that BPN-15477 would promote exon 10 exclusion in the mutated triplet. Generation and evaluation of both wild type and mutant minigenes confirmed our prediction (Fig 4f). Taken together, these results suggest that BPN-15477 and other SMCs of this class might be an effective therapy for reducing the inclusion of exon 10 in MAPT, thereby delaying or ameliorating disease symptoms in patients carrying mutations that disrupt the balance of exon 10 inclusion. Our success in validating several of our CNN model predictions in both human cells and minigenes demonstrates that deep learning methods can be effectively applied to explore new therapeutic targets for known splicing modulators, and further sets the stage for evaluating their effectiveness across a wide range of human genetic diseases.

## Discussion

The development of drugs that can increase the amount of normal transcript through modulating RNA splicing in patients is a new, precisely targeted treatment approach aimed directly at the primary molecular disease mechanism without altering the genome. The recent success of splicing modulation therapies for DMD (exondys 51) and SMA (nusinersen, risdiplam rg7800, branaplam)^42,43^ has validated the utility of splicing modification as a valuable therapeutic strategy for human disorders. Here, we have identified a novel compound, BPN-15477, that corrects splicing of *ELP1* both in a minigene system and in FD patient cell lines. We have demonstrated that BPN-15477 is a useful tool compound to study the effect of SMCs on splicing and its further optimization will lead to new therapeutic interventions for other human splicing disorders.

To determine the potential of BPN-15477 to correct splicing in other genes, we developed a machine learning approach that uses sequence signatures to predict targetable splicing defects. Our CNN model identified a total of thirty-nine 5-mer motifs important for drug response, with twelve motifs accounting for most of the BPN-15477 sensitivity when motifs are located close to the 5’ splice site. Evaluation of splice site strength in drug responsive triplets where middle exon inclusion is increased showed that these exons have weaker 5’ splice sites, a finding consistent with the previously described mechanism of action of our SMCs. We have reported that the splicing defect characteristic of FD is due to the weak definition of *ELP1* exon 20 and the kinetin-analog RECTAS was shown to promote the recognition of *ELP1* exon 20 through the recruitment of U1 snRNP at the 5’ splice site^45,68^. Our CNN model predictions, combined with these previously published observations, strongly suggest that our SMCs act by promoting recognition of weakly defined exons.

Application of our CNN model to all ClinVar pathogenic mutations that disrupt splicing identified 214 human disease-causing mutations in 155 unique genes as potential therapeutic targets of BPN-15477, proving that deep learning models are a powerful approach to explore novel therapeutic targets for drugs that modify RNA splicing. As proof of principle, we validated the treatment effect on splicing for several disease-causing mutations using patient cell lines or minigenes, and demonstrate the potential therapeutic feasibility of targeting splicing in patients with cystic fibrosis (*CFTR*), cholesterol ester storage disease (*LIPA*), Lynch syndrome (*MLH1*) and familial frontotemporal dementia (*MAPT*). These findings could have significant impact for patients carrying these mutations. For example, Wolman disease and CESD are both caused by mutations in *LIPA*^55-57^. Wolman is lethal in infancy, whereas CESD patients have some residual enzyme activity and therefore have a milder clinical course. Remarkably, patients with only 3% of the normal level of *LIPA* transcript have the much milder disease CESD, and we show that BPN-15477 increases exon 8 inclusion by 10% in a patient cell line, suggesting high potential therapeutic efficacy^58^. Similarly, our demonstration that BPN-15477 significantly increases correctly spliced CFTR transcript suggests that treatment could increase the production of functional CFTR ion channel in CF patients carrying the c.2988G>A mutation.

The ability of BPN-15477 to promote the exclusion of exon 10 in the *MAPT* gene is particularly exciting given its role in frontotemporal dementia. Many FTDP-linked *MAPT* mutations alter the splicing of tau exon 10, generally leading to an increase in exon 10 inclusion and increased expression of 4R tau isoforms, thereby disrupting the 3R (exon 10 skipped) and 4R (exon 10 retained) tau ratio^63-66^. Herein we show significant exon 10 exclusion after treatment in both the WT and in the c.1866 +16 C>T minigene, which is the most common mutation associated with FTD in humans. From a therapeutic perspective, restoring the 3R/4R tau ratio has the potential to reduce or even eliminate unbound 4R tau. The results further suggest this SMC might be beneficial for other forms of FTD caused by gain of function mutations in exon 10, such as P301L, P301S or the S305N, since treatment could reduce the level of mutated transcript.

To our knowledge, this work represents the first application of a machine learning approach to analyze the global activity of a splicing modifier compound and identify new therapeutic targets. Our successful laboratory validation of several of our CNN model predictions establishes the promise of such approaches and may presage the future advances in precision medicine offered by deep learning techniques. In this study, we provide just one of the myriad examples by which new therapeutic targets for small molecules that target splicing can be discovered and broaden their value in treating human genetic disease.

## Methods

### Preparation of BPN-15477 (2-chloro-N-(4-pyridylmethyl)-7H-pyrrolo[2,3-d]pyrimidin-4-amine)

BPN-15477 was manufactured by Albany Molecular Research Inc. (AMRI) and composition of matter is covered in International application No. PCT/US2016/013553. Synthesis and analytical data for BPN-15477 are described below. All materials used in the studies were >99% pure, as assessed by analytical methods including NMR, HPLC and LC/MS.

To a stirred suspension of 2,4-dichloro-7H-pyrrolo[2,3-d]pyrimidine (**1**, 5001 mg, 26.60 mmol, 1.000 eq.), obtained from AstaTech Inc., Bristol, PA, in 1,4-dioxane (50.0 mL) was added 4-(aminomethyl)pyridine (**2**, 3420 mg, 3.20 mL, 31.6 mmol, 1.19 eq.) followed by N,N-diisopropylethylamine (4450 mg, 6.00 mL, 34.1 mmol, 1.28 eq.) at room temperature. The reaction mixture was then heated to 90 °C and stirred at that temperature overnight.

The reaction progress was monitored by LC-MS analysis of an aliquot of the reaction mixture. After 12 h, ∼6% starting material was detected by LC-MS. The reaction was quenched by water resulting in an emulsion. The mixture was filtered through Celite and then washed well with EtOAc (3×80 mL). The organic phase was separated and the aqueous phase was extracted with EtOAc (3×40 mL). The combined organic phases were washed with brine (50 mL) and then dried over sodium sulfate.

The volatiles were removed under reduced pressure to give the crude product as dark brown solid. To the crude solid was added EtOAc (100 mL). The mixture was then heated at reflux for 15 min before it was slowly cooled to room temperature. The resulting precipitate was collected by filtration, then washed well with cold EtOAc (30 mL) followed by diethyl ether (100 mL). The solid was dried under high vacuum overnight to afford 2-chloro-N-(4-pyridylmethyl)-7H-pyrrolo[2,3-d]pyrimidin-4-amine (BPN-15477) as a light brown solid (3450 mg, 13.3 mmol, 50% yield.)

LC-MS: 0.63 min (254 nm), *m/z* 260.3, 262.3 [M+H]^+^, 258.2, 260.2 [M-H]^-^; ^1^H NMR (DMSO-*d*_*6*_) δ: 11.65-11.85 (m, 1H), 8.51 (d, J=6.0 Hz, 2H), 8.45-8.50 (m, 1H), 7.28-7.40 (m, 2H), 7.09-7.21 (m, 1H), 6.53-6.74 (m, 1H), 4.61-4.81 (m, 2H).

### Cell culture

HEK-293T (ATCC) cells were cultured in Dulbecco’s modified Eagle’s medium (11995-065, D-MEM, Gibco) supplemented with 10% fetal bovine serum (FBS, 12306C, Sigma) and 1% penicillin/streptomycin (30-009-CI, Corning).

HEK-293T, stably transfected with the expression minigenes (EMGs) for the full-length coding sequence and flanking intron sequence of the Cystic Fibrosis Transmembrane Conductance Regulator (CFTR) (courtesy of Dr. Garry R. Cutting, Johns Hopkins University school of Medicine, Baltimore, Maryland) were cultured in D-MEM supplemented with 10% FBS (FBS), 1% penicillin/streptomycin and 0.1mg/mL Hygromycin (400052-5ML, Sigma).

Patient human fibroblast GM04663 (Coriell Cell Repository) carrying the c.2204+6T>C mutation in *ELP1*, GM03111 (Coriell Cell Repository) carrying the c.894G>A mutation in *LIPA* and the human wild type fibroblasts (Coriell Cell Repository) listed in Table S1 used for RNA sequence were cultured in D-MEM supplemented with 10% FBS and 1% penicillin/streptomycin.

### Treatment of Cultured Cells

Different batches of BPN-15477 were used for dissolution into 100% DMSO to yield 40 mM stock solutions. Working solutions (10X) were prepared by dilution in 5% DMSO in phosphate-buffered saline (PBS, 10010023 ThermoFisher Scientific). The final DMSO concentration in the treated or untreated cells was 0.5%. Kinetin was purchased from Sigma (K3253).

Cells to be treated with BPN-15477 are seeded at the appropriate density in specific vessels so as to reach semiconfluency at the time of treatment. HEK293T transfected with minigenes were seeded in 6 wells and patient fibroblasts in 10 cm dishes using the described media. The following day, the media was changed with regular growth media supplemented with compound or DMSO working solutions to obtain final concentrations of 60 μM BPN-15477 and 0.05% DMSO. 60 μM BPN-15477 was chosen to guarantee the maximum effect possible on splicing (Fig1b and c). Cells were collected for RNA extraction 24 hours after compound or DMSO addition.

### Transfection

HEK293T cells were seeded in 6-well culture plates at 1.20 ×10^6^ cells/well in D-MEM,10% FBS, without antibiotics and incubated overnight to reach approximately 90% confluence the day after. Transfection was performed with FuGENE® HD Transfection Reagent (E2311, Promega) using the FuGENE-DNA ratio at 3.5:1 and following manufacturer protocol. After 4 hours of incubation at 37 °C, cells were plated at a density of 3×10^4^ cells/well in a poly-L-lysine coated 96-well plate for the dual luciferase assay or at the density of 8.5×10^5^ cells/well into 6-well plates for minigene transfection. After 16 hours incubation at 37 °C, SMCs or DMSO were added at the desired concentrations as described in the next paragraph and kept in culture for other 24 hours.

### Dual-luciferase splicing assay

Rluc-FD-Fluc plasmid used for the dual-luciferase splicing assay was derived using the *ELP1* FD minigene^44^ containing the *ELP1* genomic sequence spanning exon 19-21 inserted into spcDNA3.1/V5-His Topo (Invitrogen). Firefly luciferase (FLuc) coding sequence was inserted immediately after exon 21 and renilla luciferase (RLuc) upstream of exon 19. Characterization of the assay has shown that RLuc is expressed each time a transcript is generated from the reporter plasmid, while FLuc is only expressed if exon 20 is included in the transcript, thereby keeping FLuc in-frame. Evaluation of FLuc/RLuc expression yields the percent exon inclusion in the splicing assays^49^. To perform the dual-luciferase assay HEK-293T were transfected with the described plasmid and treated with SMCs for 24 hours as illustrated above. After treatment the cells were washed once in PBS and lysed for 25 minutes at room temperature using 50µL well of passive lysis buffer (E1941, Promega). Luciferase activity was measured as previously described^49^.

### RNA isolation and RT-PCR analysis

After treatment, cells were collected and RNA was extracted with QIAzol Lysis Reagent (79306, Qiagen) following the manufacturer’s instructions. The yields of the total RNA for each sample were determined using a Nanodrop ND-1000 spectrophotometer.

Reverse transcription was performed using 0.5-1 µg of total RNA, Random Primers (C1181, Promega), Oligo(dT)15 Primer (C1101, Promega), and Superscript III reverse transcriptase (18080093, ThermoFisher Scientific) according to the manufacturer’s protocol. cDNA was used to perform PCR reaction in a 20-25 µL volume, using GoTaq® green master mix (MT123, Promega). Primers and melting temperature (T_m_) used are described (see Supplementary Methods). To measure the splicing of the minigenes, forward and reverse primers were designed to include the TOPO/V5 plasmid vector and flanking exon sequence in order to avoid endogenous gene detection. PCR reaction was performed as follows: 32 cycles of (95°C for 30 s, Tm for 30 s, 72°C for 30 s), products were resolved on a 1.5 – 3 % agarose gel, depending on the dimension of the bands to be separated, and visualized by ethidium bromide staining.

Ratios between isoforms with included or excluded middle exon were obtained using the integrated density value (IDV) for each correspondent band, assessed using Alpha 2000TM Image Analyzer and quantified by ImageJ software. The level of exon inclusion was calculated as previously described (Cuajungco et al., 2003) as the relative density value of the band representing inclusion and expressed as a percentage. Validation of the semi-quantitative RT-PCR method for the measurement of exon inclusion has been previously performed using RT-qPCR (Cuajungco et al., 2003).

### RNASeq experiment

Six different human fibroblast cell lines from healthy individuals were obtained from Coriell Institute (Table S1) and cultured in D-MEM supplemented with 10% FBS and 1% penicillin/streptomycin. Cells were counted and plated in order to achieve semi-confluence after eight days. Twenty-four hours after plating, the medium was changed and cells were treated with BPN-15477 or DMSO to a final concentration of 30 µM and 0.5%, respectively. DMSO was used as vehicle and the concentration of BPN-15477 was chosen based on our previous studies, since at this concentration BPN-15477 induces robust splicing changes and ELP1 protein increase. After seven days of treatment, cells were collected, and RNA was extracted using the QIAzol Reagent following the manufacturer’s instructions. RNASeq libraries were prepared by the Genomic and Technology Core (GTC) at MGH using strand-specific dUTP method ^69^. In brief, RNA sample quality (based on RNA Integrity Number, or RIN) and quantity was determined using the Agilent 2200 TapeStation and between 100-1000 ng of total RNA was used for library preparation. Each RNA sample was spiked with 1 µl of diluted (1:100) External RNA Controls Consortium (ERCC) RNA Spike-In Mix (4456740, ThermoFisher Scientific) alternating between mix 1 and mix 2 for each well in the batch. Samples were then enriched for mRNA using polyA capture, followed by stranded reverse transcription and chemical shearing to make appropriate stranded cDNA inserts. Libraries were finished by adding Y-adapters, with sample specific barcodes, followed by between 10-15 rounds of PCR amplification. Libraries were evaluated for final concentration and size distribution by Agilent 2200 TapeStation and/or qPCR, using Library Quantification Kit (KK4854, Kapa Biosystems), and multiplexed by pooling equimolar amounts of each library prior to sequencing. Pooled libraries were 50 base pair paired-end sequenced on Illumina HiSeq 2500 across multiple lanes. Real time image analysis and base calling were performed on the HiSeq 2500 instrument using HiSeq Sequencing Control Software (HCS) and FASTQ files demultiplexed using CASAVA software version 1.8. RNASeq reads were mapped to the human genome Ensembl GRCh37 by STAR v2.5.2a allowing 5% mismatch^70^. Exon triplet index was built according to transcriptome Ensembl GRCh37 version 75. Reads spliced at each exon triplet splice junction was calculated by STAR on the fly.

### Differential splicing analysis

For each exon triplet in a certain biological replicate, ψ was calculated as 0.5*(R1+R2)/(0.5*(R1+R2)+R3) according to Fig.2a. The average ψ was calculated for treated and untreated condition, followed by the calculation of ψ change. For a certain exon triplet in a certain biological replicate, a 2×2 table was created, where the four cells of the table represent number of reads supporting middle exon inclusion and skipping before and after treatment. Thus, for each exon triplet, totally six 2×2 tables were created for six biological replicates. Cochran-Mantel-Haenszel test was applied to test whether there is an association between treatment and splicing across all replicates (namely whether the cross-replicate odds ratio is 1 or not). For each exon triplet, a *p* value of Cochran-Mantel-Haenszel test was reported. Benjamini-Hochberg false-discovery-rate (BH FDR) correction was finally applied to p values of all triplets.

### CNN model

Our CNN network contains two convolutional layers and one hidden layer (Supplementary Fig. 1a) and it was trained using Basset framework^41^. The training set consists of 178 inclusion-responded, 476 exclusion-responded and 268 unchanged exon triplets. The validation sets consist of 51 inclusion-responded, 136 exclusion-responded and 76 unchanged exon triplets. The test set consists of 25 inclusion-responded, 68 exclusion-responded and 38 unchanged exon triplets. The above three sets were assigned randomly in Python using seed of 122. For each exon-triplet, the sequences from UI_1_, I_1_X, XI_2_ and I_2_D, each of which consisted of 25 bp of exonic sequence and 75bp of intronic sequence, are concatenated and then one-hot coded into an input matrix with size of 4 × 400. The first round of convolution is applied with fifty 4 × 5 weight matrices, converting the input matrix into a 50 × 396 convoluted matrix, in which each row represents the convolution of one weight matrix. Then the convoluted matrix is nonlinearly transformed by rectified linear unit (ReLU) function and max pool stage takes the maximum of two adjacent positions of each row, shrinking the output matrix to a size of 50 × 198. Then the second around of convolution applies fifty 4 × 2 weight matrices, followed by the same ReLU transformation and max pool of the first round. The output is converted to 1 × 500 matrix to initiate the hidden layer, where a fully connected network is built with 90% dropout rate. The output from the hidden layer is ReLU transformed again and is then linearly transformed into a vector of three values, representing three different treatment responses. The final sigmoid nonlinearity maps each element in the vector to a value between 0 and 1, considering as the probability of drug responsiveness. In each epoch of training, average of area under curve (AUC) was measured on the validation set across the prediction of three treatment responses. The training and validation loss in terms of binary cross-entropy were measured on the training set and validation set respectively. The training process stopped if there is no improvement of average of AUC in 10 consecutive epochs. In this study, we stopped at 12^th^ epoch to avoid overfitting (Supplementary Fig. 1b).

### Examination of motif contribution

To examine motif contribution in classification, the validation set was used as model input. For each motif’s contribution to be measured, its total output of the first convolutional layer from was manually set to the mean of all output. The model was then taken forward without tuning other parameters and the new AUC of each class was calculated. The contribution of that motif was measured as the difference between the new AUC and the original AUC of each class. All motifs were investigated in this way in turn. A motif whose contribution is more than 0 in any class is considered a true identified motif. A motif whose contribution is no less than 0.1 in any class is considered a top contributor in drug response prediction.

### Analysis of positional importance

To examine each motif contribution in classification, the validation set was used as model input. For each motif whose positional importance was to be measured, the position-wise output of the first convolutional layer from that motif was manually set to the mean of all the convolutional outputs. The model was then taken forward without tuning other parameters and the new loss, measured by binary cross-entropy, of the model was calculated. The importance of that motif at that tuned position was measured as the difference between the new loss of the model and the original loss of the model. All the positions of that motif were investigated in turn. Each following motif was systematically investigated in the same fashion.

### *In silico* saturated mutagenesis

For a given input sequence, each position was *in silico* mutated to the other three alternative nucleotides. The loss of the model using mutated sequences were calculated and compared to the loss derived from the original sequence. The maximum change of loss at each position was recorded. Nucleotides were drawn proportionally to the change of loss, beyond a minimum height of 0.25.

### Standardized probability from prediction

To determine the final drug response (inclusion, exclusion or unchanged) from the prediction, the raw prediction score from the model was standardized. For each class, a cutoff representing 95% specificity of that class was identified on the validation set. The intermediate score of each class was calculated as the raw prediction score divided by the cutoff of that class. The standardized probability for each class was then calculated as the intermediate score divided by the sum of intermediate scores of three classes.

### *k-*mer enrichment analysis

The sequences at −3 to +7 bp of the 5’ splice sites of the middle exons for inclusion, exclusion and unchanged exon triplets were extracted. For each class, 5-mer enrichment was estimated against the other two classes using Discriminative Regular Expression Motif Elicitation (DREME) from THE MEME Suite with the parameter “*-p, -n, -dna -e 0.05* and *-k 5*”^71^.

### Splicing strength analysis

Splice strength was measured by maximum entropy model^53^. As described in the original study, the measurement in this paper takes the short sequence of 9bp and 23bp flanking splice junctions, depending on whether it’s 5’ or 3’ side.

### Candidate selection for minigene validation

To validate whether the CNN model correctly predicts BPN-15477 treatment response of mutated exon triplets, the following rules were applied to select suitable exon triplet: 1) exon triplets with a total length, including introns, less than 1.5 kb suitable for cloning; 2) exon triplets whose splicing changes were detectable in fibroblast RNASeq after BPN-15477 treatment, so that their drug responses could be used as positive control for the mutated minigenes; 3) The minigene recapitulates the same splicing change measured in the fibroblast by RNASeq to guarantee that the splicing process is intact in the minigene

### Minigene generation

Wild-type and mutant double-stranded DNA (dsDNA) fragments, selected based on low nucleotide length and exon-skipping probability, were ordered through GENEWIZ (FragmentGENE). Adenosine was enzymatically attached to DNA fragment 3’ ends with Taq Polymerase in the presence of 200 nM dATP and 2 mM MgCl_2_ at 70 ^o^C for 30 min. Fragments were ligated into linearized pcDNA™3.1/V5-His TOPO® TA plasmid (K480001 ThermoFisher Scientific) according to manufacturer’s instructions. After colony selection and sequence confirmation, each plasmid was finally purified using MIDIprep kit (740410, NucleoBond® Xtra Midi, Takara, Mountain View, CA). Concentrations were determined using nanodrop spectrometer.

### SpliceAI prediction on ClinVar pathogenic mutations

The VCF file recording ClinVar (version 20190325) mutations was downloaded. The pathogenic/likely pathogenic mutations were extracted and fed to SpliceAI (https://github.com/illumina/SpliceAI). In the prediction from SpliceAI, any mutation with any SpliceAI score no less than 0.2 was considered alter splicing. Therefore, such mutation, together with its influenced splice junction (ISJ) and SpliceAI score were recorded.

### Rescue definition and prediction

For an exon triplet, the coordinates of the two domains of the middle exon were compared to those ISJs discovered by SpliceAI. If either domain overlapped with an ISJ and its SpliceAI score indicated a splicing gain, the exon triplet was considered with promoted exon inclusion under that corresponding mutation. On the other hand, if the 5’ splice site of the middle exon overlapped with an ISJ and its SpliceAI score indicated a splicing loss, the exon triplet was considered as exon skipping under that corresponding mutation.

Three kinds of rescue were considered: 1) if a mutated exon triplet was predicated (by SpliceAI) to cause exon skipping and it was predicted (by our CNN model) to have an inclusion response after BPN-15477 treatment; 2) if a mutated exon triplet was predicated (by SpliceAI) to cause promoted exon inclusion and it was predicted (by our CNN model) to have an exclusion response after BPN-15477 treatment; and 3) if a mutated exon triplet generated a pre-mature termination codon (PTC) inside middle exon and it was predicted (by our CNN model) to have an exclusion response after BPN-15477 treatment and its reading frame was not shifted after skipping the middle exon.

### Allele frequency from gnomAD

VCF files for both human exome and genome sequencing were downloaded from gnomAD (v2.1.1). The corresponding ClinVar mutations were located in these VCF files via their SNP IDs. If the short variant was found only in exome or only in genome sequencing VCF, the reported minor allele frequency was then used. If short variant was found in both exome or in genome sequencing, the combined frequency was calculated as (*AC1*+*AC2*)/(*AN1*+*AN2*), where *AC1* and *AC2* are the allele counts exome and genome sequencing respectively and *AN1* and *AN2* are the total sample sizes for exome and genome sequencing respectively.

### Statistical analyses

For differential splicing analysis, Cochran-Mantel-Haenszel test was applied followed by FDR correction. An FDR < 0.1 and Δψ ≥ 0.1 was considered as inclusion-response after the treatment (Fig. 2b). Any triplet with FDR < 0.1 and Δψ ≤ −0.1 was considered as exclusion-response after the treatment (Fig. 2b). Any exon triplet, whose ψ before-treatment range from 0.1 to 0.9 and ψ change is less than 0.01, was considered as unchanged-response. For Pearson correlation (Fig. 2c), the grey zone indicates 95% confident intervals. The p values were then adjusted by Bonferroni correction. For splicing strength comparison (Fig. 3c), the maximum entropy was compared among inclusion, exclusion and unchanged groups at each splice junction, using unpaired Welch’s *t* test followed by Bonferroni correction. For the boxplots (Fig. 3c), the middle lines inside boxes indicate the medians. The lower and upper hinges correspond to the first and third quartiles. Each box extends to 1.5 times inter-quartile range (IQR) from upper and lower hinges respectively. Outliers were not shown. For RT-PCR comparison (Fig. 3-4), unpaired Student’s *t* test was applied. For the distance comparison (Fig. 4b), the Kolmogorov-Smirnov (K-S) test was applied. All the tests are done in two-tailed. In all plots, the error bars indicate the 95% confident interval if not specified. In all plots, the significance levels were marked by * *p* < 0.05, ** *p* < 0.01 and *** *p* < 0.001, where the significance levels always refer to the adjusted *p* values if the adjustment was mentioned above.

## Acknowledgements

We thank the NIH Blueprint lead development team members, Dr. Charles Cywin, Dr. Ronald Franklin, Dr. Ronald White, Jamie Driscoll, Dr. Juan Marugan, Dr. Enrique Michelotti, Dr. Amir Tamiz, Dr. Bruce Molino, Dr. Douglas Kitchen, Keith Barnes, Dr. Katrina Gwinn and Dr. Rebecca Farkas for their longstanding collaboration and helpful discussions. We thank Dr. Garry R. Cutting at Johns Hopkins University school of Medicine for kindly providing the HEK293 cells stably expressing the minigene containing the full length *CFTR* coding sequence carrying the c.2988G>A mutation (EMG-MUT). This work was supported by National Institute of Health grants (U01NS078025, R21NS095437, R01NS102423 and R37NS095640 to S.A.S. and M.E.T.).

## Author Contributions

D.G., E.M., M.S., M.E.T. and S.A.S. conceived and designed the research. D.G. performed all of the *in silico* analyses and built the CNN model for new target prediction. E.M., M.S., A.J.K, E.M.L., Y.Y. performed experiments. W.L. performed the *K*-mer enrichment analysis. D.G., E.M., M.S. analyzed data. G.J., W.P., A.R., S.E., A.C., A.D., N.A.N., C.T., K.A.E., M.W., G.K. provided advice and participated in the medicinal chemistry. D.G., E.M., M.S., M.E.T. and S.A.S. wrote the original manuscript, which was reviewed and edited by all authors. M.E.T. and S.A.S. supervised the research.

## Personal financial interests

Susan A. Slaugenhaupt is a paid consultant to PTC Therapeutics and is an inventor in Patent US9265766B2, granted to The General Hospital Corporation, entitled “Methods for Altering mRNA Splicing and Treating Familial Dysautonomia by Administering Benzyladenine,” related to use of Kinetin and in International Application Number PCT/US2016/013553, pending to The General Hospital Corporation and the United States Department of Health and Human Services, entitled “Compounds for Improving mRNA Splicing,” related to use of BPN-15477.

Wencheng Li, Amal Dakka, Nikolai A. Naryshkin, Chris Trotta, Kerstin Effenberger., Matthew Woll, Gary Karp, Yong Yu, Vijayalakshmi Gabbeta are employees of PTC Therapeutics, Inc., a biotechnology company. In connection with such employment, the author receives salary, benefits and stock-based compensation, including stock options, restricted stock, other stock-related grants, and the right to purchase discounted stock through PTC’s employee stock purchase plan.

## References

1. Cooper, T.A., Wan, L. & Dreyfuss, G. RNA and disease. Cell 136, 777–793 (2009).

2. Ritchie, D.B., Schellenberg, M.J. & MacMillan, A.M. Spliceosome structure: piece by piece. Biochim. Biophys. Acta 1789, 624–633 (2009).

3. Monani, U.R., et al. A single nucleotide difference that alters splicing patterns distinguishes the SMA gene SMN1 from the copy gene SMN2. Hum. Mol. Genet. 8, 1177–1183 (1999).

4. Cuajungco, M.P., et al. Tissue-specific reduction in splicing efficiency of IKBKAP due to the major mutation associated with familial dysautonomia. Am. J. Hum. Genet. 72, 749–758 (2003).

5. Flanigan, K.M., et al. Mutational spectrum of DMD mutations in dystrophinopathy patients: application of modern diagnostic techniques to a large cohort. Hum. Mutat. 30, 1657–1666 (2009).

6. Juan-Mateu, J., et al. Interplay between DMD point mutations and splicing signals in Dystrophinopathy phenotypes. PLoS One 8, e59916 (2013).

7. Thornton, C.A. Myotonic dystrophy. Neurol. Clin. 32, 705-719, viii (2014).

8. Pros, E., et al. NF1 mutation rather than individual genetic variability is the main determinant of the NF1-transcriptional profile of mutations affecting splicing. Hum. Mutat. 27, 1104–1114 (2006).

9. Bottillo, I., et al. Functional analysis of splicing mutations in exon 7 of NF1 gene. BMC Med. Genet. 8, 4 (2007).

10. Tzetis, M., Efthymiadou, A., Doudounakis, S. & Kanavakis, E. Qualitative and quantitative analysis of mRNA associated with four putative splicing mutations (621+3A-->G, 2751+2T-->A, 296+1G-->C, 1717-9T-->C-D565G) and one nonsense mutation (E822X) in the CFTR gene. Hum. Genet. 109, 592–601 (2001).

11. Cabello, G.M., Cabello, E.J., Jr., Fernande, O. & Harris, A. The 3120 +1G-->A splicing mutation in CFTR is common in Brazilian cystic fibrosis patients. Hum. Biol. 73, 403–409 (2001).

12. Giorgi, G., et al. Validation of CFTR intronic variants identified during cystic fibrosis population screening by a minigene splicing assay. Clin. Chem. Lab. Med. 53, 1719–1723 (2015).

13. Goina, E., Fernandez-Alanis, E. & Pagani, F. Approaches to study CFTR pre-mRNA splicing defects. Methods Mol. Biol. 741, 155–169 (2011).

14. Rhine, C.L., et al. Hereditary cancer genes are highly susceptible to splicing mutations. PLoS Genet 14, e1007231 (2018).

15. Srebrow, A. & Kornblihtt, A.R. The connection between splicing and cancer. J. Cell Sci. 119, 2635–2641 (2006).

16. Skotheim, R.I. & Nees, M. Alternative splicing in cancer: noise, functional, or systematic? Int. J. Biochem. Cell Biol. 39, 1432–1449 (2007).

17. Dlamini, Z., Mokoena, F. & Hull, R. Abnormalities in alternative splicing in diabetes: therapeutic targets. J. Mol. Endocrinol. 59, R93–R107 (2017).

18. Juan-Mateu, J., Villate, O. & Eizirik, D.L. MECHANISMS IN ENDOCRINOLOGY: Alternative splicing: the new frontier in diabetes research. Eur. J. Endocrinol. 174, R225–238 (2016).

19. Bamshad, M.J., et al. The Centers for Mendelian Genomics: a new large-scale initiative to identify the genes underlying rare Mendelian conditions. Am. J. Med. Genet. A 158A, 1523–1525 (2012).

20. Sanders, S.J., et al. Insights into Autism Spectrum Disorder Genomic Architecture and Biology from 71 Risk Loci. Neuron 87, 1215–1233 (2015).

21. Fresard, L., et al. Identification of rare-disease genes using blood transcriptome sequencing and large control cohorts. Nat. Med. 25, 911–919 (2019).

22. Landrum, M.J., et al. ClinVar: public archive of relationships among sequence variation and human phenotype. Nucleic Acids Res. 42, D980–985 (2014).

23. Jaganathan, K., et al. Predicting Splicing from Primary Sequence with Deep Learning. Cell 176, 535–548 e524 (2019).

24. Johnson, N.T., Dhroso, A., Hughes, K.J. & Korkin, D. Biological classification with RNA-seq data: Can alternatively spliced transcript expression enhance machine learning classifiers? RNA 24, 1119–1132 (2018).

25. Paggi, J.M. & Bejerano, G. A sequence-based, deep learning model accurately predicts RNA splicing branchpoints. RNA 24, 1647–1658 (2018).

26. Wang, J. & Wang, L. Deep Learning of the Back-splicing Code for Circular RNA Formation. Bioinformatics (2019).

27. Palacino, J., et al. SMN2 splice modulators enhance U1-pre-mRNA association and rescue SMA mice. Nat. Chem. Biol. 11, 511–517 (2015).

28. Hua, Y., et al. Antisense correction of SMN2 splicing in the CNS rescues necrosis in a type III SMA mouse model. Genes Dev. 24, 1634–1644 (2010).

29. Passini, M.A., et al. Antisense oligonucleotides delivered to the mouse CNS ameliorate symptoms of severe spinal muscular atrophy. Sci. Transl. Med. 3, 72ra18 (2011).

30. Aartsma-Rus, A. & van Ommen, G.J. Antisense-mediated exon skipping: a versatile tool with therapeutic and research applications. RNA 13, 1609–1624 (2007).

31. Dal Mas, A., Rogalska, M.E., Bussani, E. & Pagani, F. Improvement of SMN2 pre-mRNA processing mediated by exon-specific U1 small nuclear RNA. Am. J. Hum. Genet. 96, 93–103 (2015).

32. Havens, M.A., Duelli, D.M. & Hastings, M.L. Targeting RNA splicing for disease therapy. Wiley interdisciplinary reviews. RNA 4, 247–266 (2013).

33. Sinha, R., et al. Antisense oligonucleotides correct the familial dysautonomia splicing defect in IKBKAP transgenic mice. Nucleic Acids Res. 46, 4833–4844 (2018).

34. Vigevani, L. & Valcarcel, J. Molecular biology. A splicing magic bullet. Science 345, 624–625.

35. Naryshkin, N.A., et al. Motor neuron disease. SMN2 splicing modifiers improve motor function and longevity in mice with spinal muscular atrophy. Science 345, 688–693.

36. Woll, M.G., et al. Discovery and Optimization of Small Molecule Splicing Modifiers of Survival Motor Neuron 2 as a Treatment for Spinal Muscular Atrophy. J. Med. Chem. 59, 6070–6085 (2016).

37. Ratni, H., et al. Discovery of Risdiplam, a Selective Survival of Motor Neuron-2 (SMN2) Gene Splicing Modifier for the Treatment of Spinal Muscular Atrophy (SMA). J. Med. Chem. 61, 6501–6517 (2018).

38. Esteva, A., et al. A guide to deep learning in healthcare. Nat. Med. 25, 24–29 (2019).

39. Ozaki, K., et al. Functional SNPs in the lymphotoxin-alpha gene that are associated with susceptibility to myocardial infarction. Nature genetics 32, 650–654 (2002).

40. Golub, T.R., et al. Molecular classification of cancer: class discovery and class prediction by gene expression monitoring. Science 286, 531–537 (1999).

41. Kelley, D.R., Snoek, J. & Rinn, J.L. Basset: learning the regulatory code of the accessible genome with deep convolutional neural networks. Genome Res. 26, 990–999 (2016).

42. Chiriboga, C.A., et al. Results from a phase 1 study of nusinersen (ISIS-SMN(Rx)) in children with spinal muscular atrophy. Neurology 86, 890–897 (2016).

43. Cheung, A.K., et al. Discovery of Small Molecule Splicing Modulators of Survival Motor Neuron-2 (SMN2) for the Treatment of Spinal Muscular Atrophy (SMA). J. Med. Chem. 61, 11021–11036 (2018).

44. Slaugenhaupt, S.A., et al. Rescue of a human mRNA splicing defect by the plant cytokinin kinetin. Hum. Mol. Genet. 13, 429–436 (2004).

45. Yoshida, M., et al. Rectifier of aberrant mRNA splicing recovers tRNA modification in familial dysautonomia. Proc. Natl. Acad. Sci. U. S. A. 112, 2764–2769 (2015).

46. Hims, M.M., et al. Therapeutic potential and mechanism of kinetin as a treatment for the human splicing disease familial dysautonomia. J. Mol. Med. (Berl.) 85, 149–161 (2007).

47. Axelrod, F.B., et al. Kinetin improves IKBKAP mRNA splicing in patients with familial dysautonomia. Pediatr. Res. 70, 480–483 (2011).

48. Gold-von Simson, G., et al. Kinetin in familial dysautonomia carriers: implications for a new therapeutic strategy targeting mRNA splicing. Pediatr. Res. 65, 341–346 (2009).

49. Salani, M., et al. Development of a Screening Platform to Identify Small Molecules That Modify ELP1 Pre-mRNA Splicing in Familial Dysautonomia. SLAS Discov, 2472555218792264 (2018).

50. Wang, E.T., et al. Alternative isoform regulation in human tissue transcriptomes. Nature 456, 470–476 (2008).

51. Sakuma, M., Iida, K. & Hagiwara, M. Deciphering targeting rules of splicing modulator compounds: case of TG003. BMC Mol. Biol. 16, 16 (2015).

52. Chiara, M.D., et al. Identification of proteins that interact with exon sequences, splice sites, and the branchpoint sequence during each stage of spliceosome assembly. Mol. Cell. Biol. 16, 3317–3326 (1996).

53. Yeo, G. & Burge, C.B. Maximum entropy modeling of short sequence motifs with applications to RNA splicing signals. J. Comput. Biol. 11, 377–394 (2004).

54. Bowden, K.L., et al. Lysosomal acid lipase deficiency impairs regulation of ABCA1 gene and formation of high density lipoproteins in cholesteryl ester storage disease. J. Biol. Chem. 286, 30624–30635 (2011).

55. Reiner, Z., et al. Lysosomal acid lipase deficiency--an under-recognized cause of dyslipidaemia and liver dysfunction. Atherosclerosis 235, 21–30 (2014).

56. Saito, S., Ohno, K., Suzuki, T. & Sakuraba, H. Structural bases of Wolman disease and cholesteryl ester storage disease. Mol. Genet. Metab. 105, 244–248 (2012).

57. Zhang, B. & Porto, A.F. Cholesteryl ester storage disease: protean presentations of lysosomal acid lipase deficiency. J. Pediatr. Gastroenterol. Nutr. 56, 682–685 (2013).

58. Aslanidis, C., et al. Genetic and biochemical evidence that CESD and Wolman disease are distinguished by residual lysosomal acid lipase activity. Genomics 33, 85–93 (1996).

59. Scott, S.A., et al. Frequency of the cholesteryl ester storage disease common LIPA E8SJM mutation (c.894G>A) in various racial and ethnic groups. Hepatology 58, 958–965 (2013).

60. Sharma, N., et al. Experimental assessment of splicing variants using expression minigenes and comparison with in silico predictions. Hum. Mutat. 35, 1249–1259 (2014).

61. Heaney, D.L., Flume, P., Hamilton, L., Lyon, E. & Wolff, D.J. Detection of an apparent homozygous 3120G>A cystic fibrosis mutation on a routine carrier screen. J. Mol. Diagn. 8, 137–140 (2006).

62. Pande, M., et al. Cancer spectrum in DNA mismatch repair gene mutation carriers: results from a hospital based Lynch syndrome registry. Fam. Cancer 11, 441–447 (2012).

63. Hutton, M., et al. Association of missense and 5’-splice-site mutations in tau with the inherited dementia FTDP-17. Nature 393, 702–705 (1998).

64. Spillantini, M.G., et al. Mutation in the tau gene in familial multiple system tauopathy with presenile dementia. Proc. Natl. Acad. Sci. U. S. A. 95, 7737–7741 (1998).

65. Hong, M., et al. Mutation-specific functional impairments in distinct tau isoforms of hereditary FTDP-17. Science 282, 1914–1917 (1998).

66. Connell, J.W., et al. Quantitative analysis of tau isoform transcripts in sporadic tauopathies. Brain Res. Mol. Brain Res. 137, 104–109 (2005).

67. Neumann, M., et al. A new family with frontotemporal dementia with intronic 10+3 splice site mutation in the tau gene: neuropathology and molecular effects. Neuropathol. Appl. Neurobiol. 31, 362–373 (2005).

68. Ibrahim, E.C., et al. Weak definition of IKBKAP exon 20 leads to aberrant splicing in familial dysautonomia. Hum. Mutat. 28, 41–53 (2007).

69. Jiang, L., et al. Synthetic spike-in standards for RNA-seq experiments. Genome Res. 21, 1543–1551 (2011).

70. Dobin, A., et al. STAR: ultrafast universal RNA-seq aligner. Bioinformatics 29, 15–21 (2013).

71. Bailey, T.L. DREME: motif discovery in transcription factor ChIP-seq data. Bioinformatics 27, 1653–1659 (2011).

