## Supplementary Information and Figures for "A deep learning approach to identify new gene targets of a novel therapeutic for human splicing disorders"

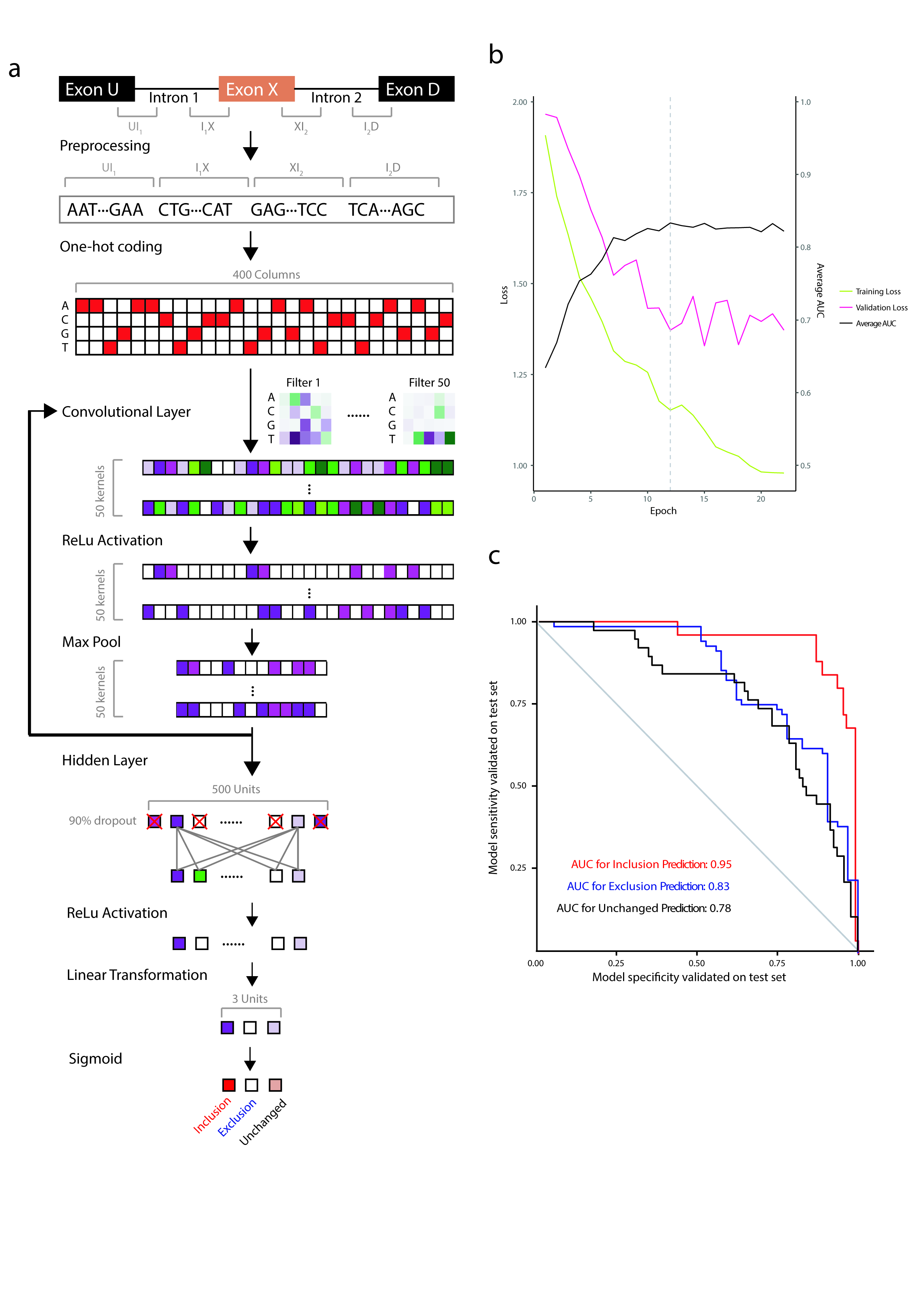

**Supplementary Fig. 1 Training process of CNN**

**(a)** CNN model work flow. For each exon-triplet, the sequences from UI_1_, I_1_X, XI_2_ and I_2_D are concatenated and then one-hot coded. In the model, two rounds of convolution were applied before the hidden layer. Each round of convolution consists of a convolution layer of fifty filters, a ReLU activation layer and a max pooling layer of size 2. After two rounds of convolution, the output is converted and connected to a hidden layer with 90% dropout rate. The output from the hidden layer is ReLU transformed again and is then linearly transformed into a vector of three, representing three different treatment responses. The final nonlinear sigmoid maps each element in the vector to a value between 0 and 1, and it is considered as the probability of drug responsiveness. **(b)** Training progress of the CNN model. The x axis represents the number of epochs iterated during training. The left y axis shows loss score measured by binary entropy while the right y axis shows the average AUC of prediction from three classes. The red line and blue line show training and validation loss respectively along the growth of epochs. The black line shows the improvement of AUC along the growth of epochs. The vertical dashed line indicates the stop of training to avoid overfitting. **(c)** AUC curves of prediction on each class using the test set. The x axis represents specificity while the y axis represents sensitivity. The solid grey line indicates the boundary beneath which the prediction is not better than a random guess. The AUC for inclusion, exclusion and unchanged response are marked in red, blue and black, respectively.

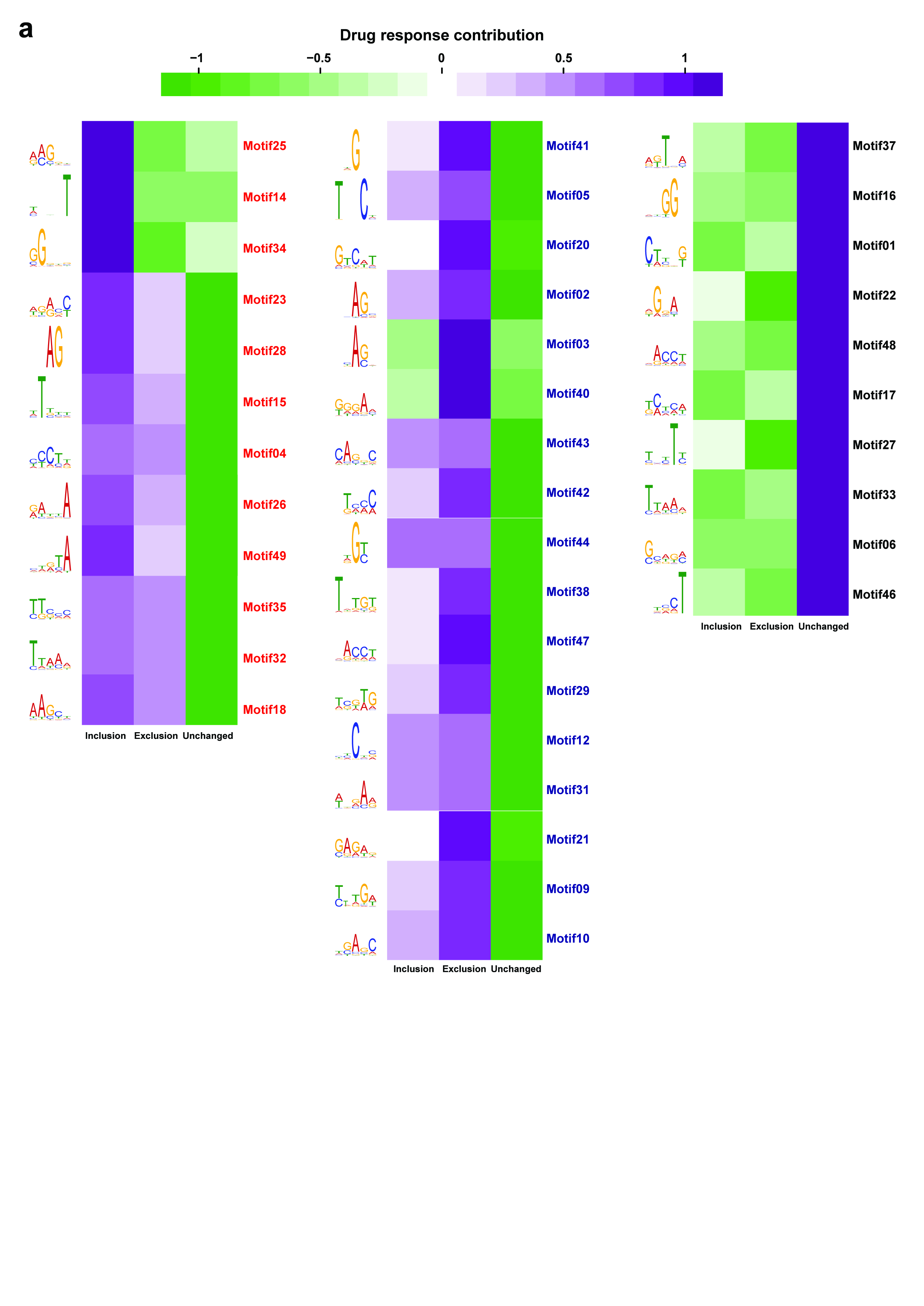
**Supplementary Fig. 2 Motifs identified by the CNN model**

**(a)** Heatmap shows all motifs identified by the model. The color indicates the directional contribution of each motif. The purple domain indicates positive contribution while the green domain indicates negative contribution. The LOGO plot of each motif is shown on the left side of the heatmap. The motif names are displayed on the right side of the heatmap. Colors assigned to the motif names, suggest their contribution to inclusion (red), exclusion (blue) and unchanged (black) response respectively.

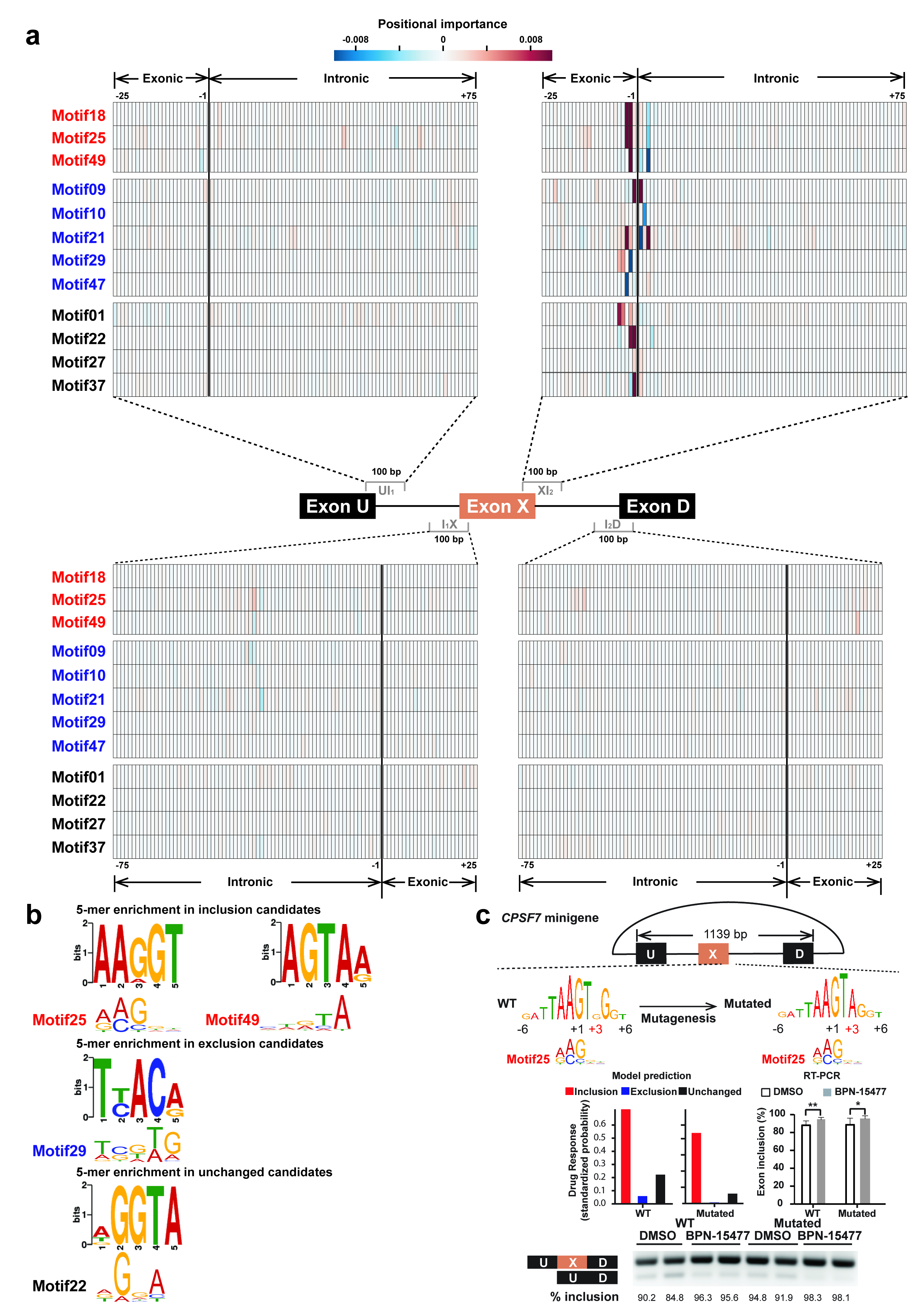
**Supplementary Fig. 3 Positional importance of motifs**

**(a)** Heatmap of motif importance at each 100bp exon boundary. Each thick vertical line shows the exon-intron boundary. Only the top twelve motifs are shown according to Fig. 3a. The motifs are grouped and colored based on their classification contribution according to Fig. 3a, representing inclusion (red), exclusion (blue) and unchanged (black) response, respectively. **(b)** 5-mer enrichment analysis at -3 to +7bp of the 5’ splice sites of the middle exons for inclusion, exclusion and unchanged classes. Their closely related CNN motifs are indicated beneath. **(c)** *Upper row*: Summary of minigene generation for *CPSF7.* The length of the exon triplet cloned into minigene is shown. The sequences around of the 5’ splice sites of the middle exons are shown in LOGO plots. The red coordinate number indicates the position of mutation relative to the 5’ splice site. Their closely matched CNN motifs are indicated beneath. *Middle row*: Splicing changes of the middle exon in both wildtype and mutated exon triplets, predicted by the CNN model (*left*) and measured by RT-PCR of the minigene (*right*). The RT-PCR was done in triplicates to make the bar plot (two-tailed, unpaired Student’s *t* test). *Bottom* *row:* Example of splicing changes induced by the treatment using a minigene splicing assay. The percentage of the middle exon inclusion are indicated beneath each gel, from treated and untreated conditions in both wildtype and mutated minigenes. Each of the gels is one of the triplicates used to generate the bar plot in *middle row* respectively. The upper bands indicate the isoforms in which the middle exon is included while the lower bands indicate the isoforms in which the middle exon is skipped. The statistical significance is determined via two-tailed, unpaired Student’s test. In the figure, **p* < 0.05; ***p* < 0.01; ****p*<0.001.

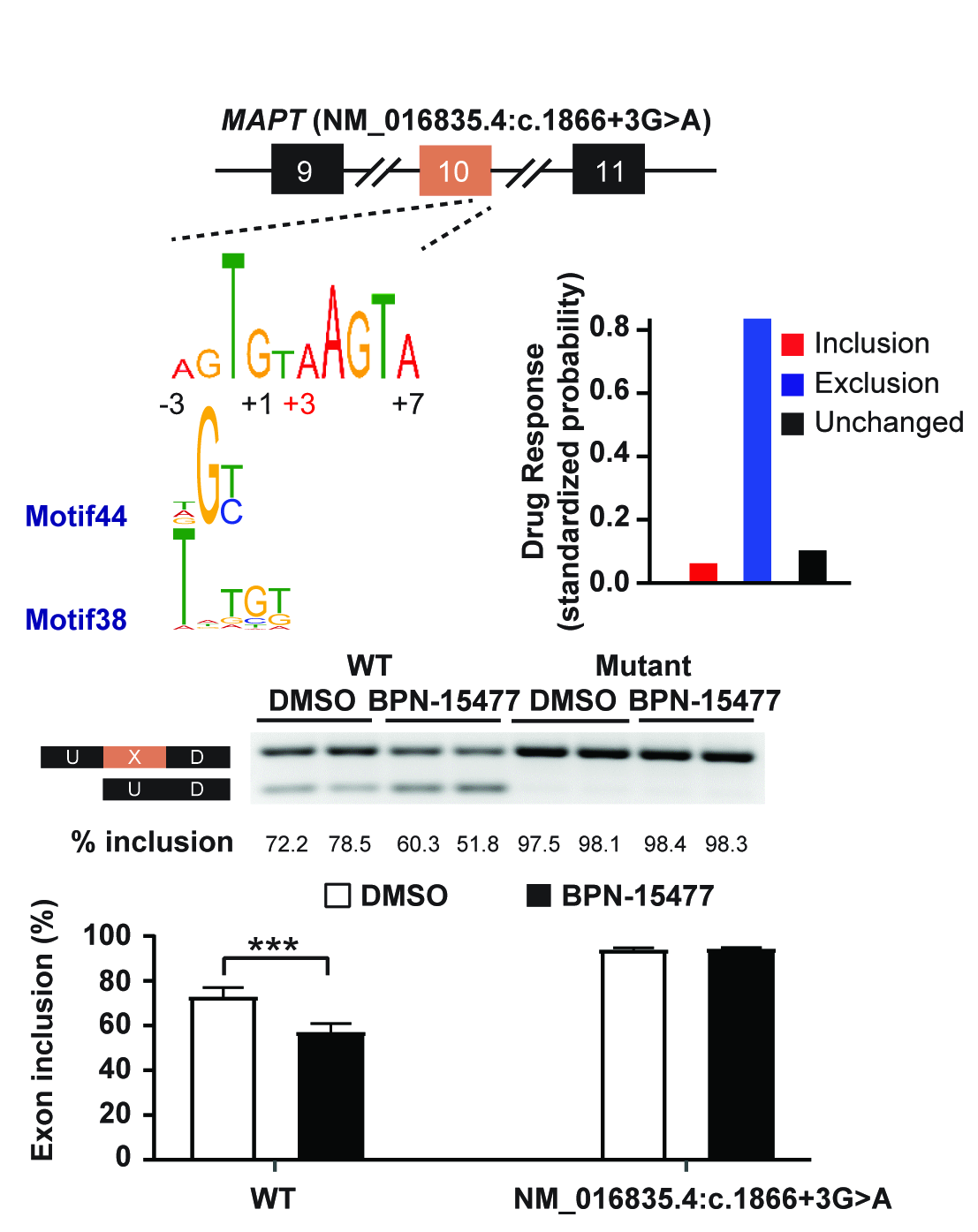

**Supplementary Fig. 4 Treatment effect on *MAPT* exon 10 splicing in the c.1866 +3 G>A mutated minigene**

*Upper row*: Schematic of the MAPT minigene construct. The sequence around the 5’ splice site of the middle exon is shown in LOGO plots, with closely matched CNN motifs indicated below. The red coordinate number indicates the position of the mutation relative to the 5’ splice site. The bar plots show that the CNN model predicts increased exon exclusion. *Middle row*: RT-PCR validation of treatment responses in cell lines expressing the mutated minigene. The gel shows one of the triplicates used to generate the bar plots beneath. *Bottom row*: The bar plots demonstrate the splicing change promoted by BPN-15477 treatment. The statistical significance is determined via two-tailed, unpaired Student’s test. In the figure, **p* < 0.05; ***p* < 0.01; ****p*<0.001.

**Supplementary Table 1:** Wildtype (WT) human fibroblast cell lines used in the RNAseq experiment.

| **Coriell #** | **Genotype** | **Sex** | **Age** | **Race** |
| --- | --- | --- | --- | --- |
| AG16409 | WT | Male | 12 years | Caucasian |
| GM03348 | WT | Male | 10 years | Caucasian |
| GM08402 | WT | Male | 32 years | Caucasian |
| GM01652 | WT | Female | 11 years | Caucasian |
| GM02036 | WT | Female | 11 years | Caucasian |
| GM00041 | WT | Female | 3 months | Caucasian |

**Supplemetary Methods**

**Primers and melting temperature (T_m_) used for RT-PCR analysis**

| Gene | Forward 5'-3' | Reverse 5'-3' | Tm |
| --- | --- | --- | --- |
| ELP1 | CCTGAGCAGCAATCATGTG | TACATGGTCTTCGTGACATC | 58 C° |
| KLC1 | CGC AGT GGT TCC TTT AGC | CAC TGC TGC TGC TGT CG | 60 C° |
| LPIN1 | GCT GTG ATT TAC CCT CAG TCA GC | CTT AGC AGC CTG CGG CAG C | 64 C° |
| HSD17B4 | GCA GAA AGA GGA GCG TTA | GTT GGC CAC TGC TTT TCC | 56 C° |
| SLC4A7 | GCT ACA GAG GAC TGG ACG | CTA GAA CTG GAC CTG TGC TCC | 60 C° |
| LRRC28 | GAT ATA GTG CTG CAG CGT GC | CAA CTA TGT TAT TTG AGT GCA GG | 60 C° |
| AFMID | GCC TTT CTT CCT GTT CTT TCA CG | GGT GAG CAC GTT GTC CTT CT | 60 C° |
| CD99P1 | CGA CCC AGC ACC TCT TAA TTC | CGG TGG AAT CAG GCT GCT TG | 62 C° |
| CRYZ | GCA CTG CTG GTA CTG AGG AAG | CTT TGC CAT GGT GTC TCG TGG | 64 C° |
| IP6K2 | AAC AAG CCA AGG AGC CAA GA | ATT CAG GCC ACA CTT CCC TG | 62 C° |
| EVC | TGC CCT GAA GCT GAT GAA GG | GGT GCC AGC GTC TGC TTC | 62 C° |
| EPB41L2 | GGA GAA GTA CCT GAT GCC GAC | CTC ACT CTC ACT GCT GCT G | 60 C° |
| CYLD | GAT GGT TCT ACA CAG CCA CC | CTT CCC AGT AGG GTG AAG TGA C | 62 C° |
| MEGF6 | CTGGTTTGGAGAGGCCTGTG | GGGACGGACTGCAACCTCA | 60 C° |
| MYEF2 | GTACCGTGGTGCGATGACTA | TCTGACAAATATCTGGTTGCCT | 60 C° |
| ASXL1 | GCCTCGAGTTGTCCTGACTC | TTCAGGCAGGAGGAAGAGGA | 60 C° |
| SPTAN1 | CGATCGTCAGGGTTTTGTGC | TTCTGGAGCACCTCAACCTG | 60 C° |
| COPS8 | CTGAGGGACAGTCTGGGGTT | ATGGAGCAAATATAAAGCTAGAAGC | 60 C° |
| PDZD11 | GTCAGTGAGCGGAGTCTGAG | TGAGGAGGAATCCATGCTGG | 60 C° |
| KTN1 | AGGCAGAGATGGAACGATCT | TGCAAATCACCAGCTACCTTCT | 60 C° |
| SPPL2A | TCATGGTTGAACTCGCAGCT | TGAGGCACACACTCATTACTGA | 60 C° |
| LIPA | CCCAGAGTGCGTTTTTGAA | CCCAGTCAAAGGCTTGAAAC | 60 C° |
| CFTR | GCAGTGATTATCACCAGC | GGAGGAAATATGCTCTCAAC | 58 C° |
| PTEN | TTGCACAATATCCTTTTGAAGACCA | TTAGCATCTTGTTCTGTTTGTGGA | 60 C° |
| CPSF7 | ATTGCCCTTGACCCAGAGTT | GAGCGAATAACCTGGATCAGC | 60 C° |
| SETD5 | TGTGGTGGAATTGCCCTTAC | GCAGAATTGCCCTTCTGATA | 60 C° |
| MLH1 | GTGGAATTGCCCTTGAGC | CACTGTGCTGGATATCTGCTG | 60 C° |
| PARP6 | GGAATTGCCCTTGATCATCT | AATTGCCCTTCTGCAGTTTG | 60 C° |
| MAPT | ACCCAAGTCGCCGTCTTCCGCC | CACCTTGCTCAGGTCAACTGGT | 60 C° |
